# Assessment of inter-individual variation in metabolism of flavonoids from bilberry and grape seed extracts using an *in vitro* digestion and faecal fermentation model

**DOI:** 10.64898/2026.03.02.709000

**Authors:** T. Grohmann, P. A. Kroon, M. Philo, G. Horgan, X. Zhang, U. Balaseviciute, A. W. Walker, W. R. Russell, N. Hoggard, B. de Roos

**Affiliations:** Rowett Institute, University of Aberdeen, Foresterhill, Ashgrove Road West, AB25 2ZD, Aberdeen, Scotland; Quadram Institute Bioscience, Rosalind Franklin Road, Norwich Research Park, NR4 7UQ, Norwich, United Kingdom; Biomathematics & Statistics Scotland, Aberdeen, Scotland; By-Health Ltd Co, No.3 Kehui 3rd Street, No.99 Kexue Avenue Central, Luogang District, Guangzhou, China

**Keywords:** *in vitro* digestion, bilberry, grape seed, extracts, flavonoids, faecal microbiota

## Abstract

The gut microbiota plays an essential role in the conversion of anthocyanins and (epi-)catechins into smaller phenolic acids, which are then absorbed into the blood stream.

The phenolic composition of a commercial bilberry extract and grape seed extract was assessed, as well as a formulation extract containing a combination of both extracts. The extracts were subjected to an *in vitro* salivary, gastric and intestinal digestion environment, based on the INFOGEST Model. The solid fraction end-product of the combined extract from the *in vitro* digestion was further fermented with faecal samples from six healthy donors, for 72 hours, to assess phenolic acid metabolism, short-chain fatty acid formation and changes in microbial composition.

During the *in vitro* digestion, flavonoid content in all three extract samples (bilberry, grape seed and the formulation extracts) decreased significantly. In the process of anthocyanin and flavonoid digestion, smaller phenolic acid compounds such as benzoic acid, cinnamic acid and mandelic acid increased in bilberry, grape seed and formulation extract samples. All faecal donors harboured unique microbiota compositions, however all faecal microbiota were able to fully convert catechin/epicatechin, the most prominent flavonoids in the formulation extract sample, into smaller phenolic metabolites (phenylacetic, phenylpropionic and benzoic acids) within 24 hours. Using 16S rRNA gene amplicon sequencing, *Anaerobutyricum* and *Enterocloster* spp. were correlated with catechin/epicatechin metabolism in the fermentation procedure, however, in single bacterial strain fermentation experiments with the formulation extract or catechin standard, these bacteria were not capable of metabolising flavonoids.

**Highlights:**

- Faecal microbiota converted (epi-)catechin to phenolic metabolites within 24 h.
- (Epi-)catechin correlated negatively with *Anaerobutyricum* and *Enterocloster* spp.
- Faecal bacterial cultures did not show (epi-)catechin metabolism capacity.

## 1 Introduction

Consumption of fresh berries, tea and coffee, as well as their extracts, has been shown to beneficially impact on blood glucose and cholesterol metabolism, and blood pressure, in some human intervention studies (Giacco et al., 2020; Grohmann et al., 2023; Williamson et al., 2018). Berries are high in flavonoids such as anthocyanins and procyanidins (Williamson & Manach, 2005) that occur primarily in glycated and polymeric forms (Kelm et al., 2004; Shehata et al., 2023). Their large complex structures result in a poor bioavailability in the human body, and a requirement for microbial action in the gut to transform such complex structures into more bioavailable moieties (Di Pede et al., 2022). Certain anthocyanin glycosides, dimeric (epi-)catechins, and aglycones, are unstable in the pH-neutral environment of the small intestine and are believed to spontaneously decay into smaller metabolites without enzyme or bacterial action (Shehata et al., 2023). A fraction of these smaller metabolites are absorbed in the small intestine and conjugated (glucuronidated, sulphated or methylated) in the enterocyte cells before entering the blood stream or returning to the small intestine (Kay et al., 2009; Williamson et al., 2018). Conjugated flavonoids can enter the colon through the enterohepatic circulation to be further metabolised by the gut microbiota (Valdés et al., 2015). Irrespective of the metabolic route, all metabolites are transported to the liver (Williamson et al., 2018), where further oxidation/reduction reactions and enzyme-mediated conjugation occurs via cytochrome P450 monooxygenases (CYP), flavin-containing monooxygenases (FMO) or hydrolase enzymes (Hodgson & Levi, 2004; Manach et al., 2017). It has been proposed that smaller metabolites may be more bioactive than their precursor flavonoids, potentially due to a simpler molecular structure and their prolonged transit time in the blood circulation (Fraga, 2009; Kelm et al., 2004; Pérez-Jiménez et al., 2010). Differences in the activity of metabolic enzymes, and in the diversity and activity of gut bacteria, may underpin the large inter-individual variability observed previously in responses to plant flavonoids in dietary intervention studies (Duncan et al., 2023; Manach et al., 2017). Indeed, every individual has a unique combination of resident gut microbe species (Valdes et al., 2018), which might result in a varied production of beneficial metabolites following dietary interventions (Ou & Gu, 2014). In this study, the conversion of flavonoids derived from a commercial bilberry extract, a commercial grape seed extract, and a formulation extract containing a combination of both, was studied. An *in vitro* digestion model was applied, followed by a faecal fermentation study with stool samples from six healthy donors, in order to understand how flavonoids are metabolised, and how inter-individual variation in gut microbiota composition may affect the formation of smaller phenolic metabolites. These factors could all potentially contribute to the health benefits of bilberry and grape seed extracts. Finally, individual faecal bacterial strains were fermented with the formulation extract and catechin standard to investigate their involvement in flavonoid metabolism.

## 2 Materials and methods

### 2.1 Ethics

The research was carried out in accordance with the Declaration of Helsinki. Approval from the Rowett Institute’s Ethical Review Committee was obtained prior to the start of the study. Informed consent was obtained from all subjects for the collection of faecal samples. All six faecal sample donors had not taken any antibiotics in the previous six months. The donors were requested to abstain from consuming supplements/prebiotics or probiotics for one month prior to sample collection. Faecal donors had different cultural backgrounds (Greece, Scotland, Australia, Malaysia and Turkey) and had lived in Scotland for at least one year.

### 2.2 Extracts

Commercial bilberry (Mirtoselect^®^) and grape seed (Enovita^®^) extracts were obtained from Indena, Italy. For the *in vitro* studies we used the individual extracts and a combination of both (54% (w/w) grape seed + 46% (w/w) bilberry extract), which was termed “formulation extract”.

### 2.3 *In vitro* digestion studies

Simulated digestion of extracts was performed as described previously by Minekus et al. (2014). We used 5g of the bilberry/grape seed/formulation extracts, and subjected these to simulated salivary, gastric and intestinal digestion in the presence of enzymes and under representative pH conditions. Each digestion procedure was performed twice, with one sample set terminating at the end of the gastric digestion phase, and the second sample terminating at the end of the intestinal digestion phase, so that the impact of pH conditions on the concentration of phenolic metabolites could be compared. Pellets and freeze-dried supernatants obtained after the digestion phases were stored at −70 °C until analysis, and the pellets were also subsequently used for faecal fermentation studies.

### 2.4 Faecal fermentation studies

The pellets of the formulation extracts, from the preceding *in vitro* intestinal digestion phase, and faecal samples from six healthy donors, were inoculated in triplicate in fermentation medium and cultured over a 72h period. The fermentation medium has been described previously by Duncan et al. (2016). The concentration of extract in the culture medium was calculated on the basis of an extract dose of 550mg and assuming a stomach volume of 1 L (Cueva et al., 2013; Tzounis et al., 2011), thus 27.5mg of formulation extract were added per 50 mL of fermentation medium. We ran two sets of controls, including faecal samples that were inoculated into medium without added extracts, and fermentation medium with the formulation extract without the addition of faecal samples.

The faecal samples were processed immediately into a faecal slurry (2g faecal sample blended with 8 mL sterile 30% glycerol-PBS in the gentleMACS^™^ Dissociator, Miltenyi Biotec), and 0.5mL slurry was added to culture flasks containing 50 mL fermentation medium, with or without addition of 27.5mg of formulation extract, while flushing with CO_2_ gas. The bottles were swirled and the first sample (0h) was drawn with a sterile syringe (BD Plastipak^™^, and BD Microlance^™^). The bottles were then sealed with rubber/metal caps and placed in a thermo-shaker at 37 °C. Further samples were then drawn through the rubber cap using a sterile syringe.

Samples from the fermentation were taken at t = 0h, 2h, 5h, 24h and 72h and analysed for the bioconversion of flavonoids into smaller metabolites, pH changes, the formation of short-chain fatty acids (at t = 0h, 24h, 48h and 72h only), and the composition of faecal microbiota using 16S rRNA gene amplicon sequencing (at t = 0h, 5h, 24h and 72h only).

Upon sampling, 0.1 mL aliquots were taken, and immediately mixed with 0.4 mL of internal standard for the analysis of flavonoids and metabolites (Multari et al., 2016). The remainder of the fermentation study samples were frozen at −70 °C for later analyses.

### 2.5 Profiling faecal microbiota using 16S rRNA gene amplicon sequencing

#### 2.5.1 DNA extraction and PCR

Microbial DNA extraction using the FastDNA SPIN Kit for Soil (MP Biomedicals, Germany), and PCR targeting the V1 to V2 region of the 16S rRNA gene was performed according to Chung et al. (2016). Purified DNA samples were further processed via Bead Clean-up and prepared for Illumina MiSeq sequencing (2 × 300 bp paired end read sequencing) at the Centre for Genome-Enabled Biology and Medicine (CGEBM), University of Aberdeen.

#### 2.5.2 Sequence analysis

The sequence data were analysed with the mothur software package (Schloss et al., 2009), following the data processing steps described previously by Chung et al. (2020). Data subsam pling was performed at 9100 sequences to obtain equal sequence depths for sample comparison. Clustering patterns were analysed via Bray-Curtis-based dendrograms visualised using iTOL software (Letunic & Bork, 2019).

#### 2.5.3 Short-Chain Fatty Acid (SCFA) analysis

The SCFA extraction and analysis was performed according to the protocol described previously by Richardson et al. (1989). Derivatised SCFA samples were measured on an Agilent HP 6890 GC system using an Agilent 190912-333 HP-1: Methyl Siloxane column.

### 2.6 Targeted analysis of flavonoids and phenolic metabolites

Flavonoid extraction was performed in a three-stage process as previously described by Russell et al. (2011).

### 2.7 Targeted analysis of anthocyanins

The analysis of anthocyanin content in the extracts and the *in vitro* digested extract samples was performed at the Quadram Institute Bioscience (Norwich Research Park, UK). Extract powders (20mg) were weighed into 1.5 mL tubes, mixed with extraction solvent (1 mL, formic acid : methanol : water 1:14:35), vortexed for 10 min and centrifuged at 13,000 rpm for 10 min. The supernatant was filtered (0.45 µm) and analysed with Ultra-Performance Liquid Chromatography (UPLC-MS) in a C_18_ HSS T3 100 mm × 2.1 mm × 0.017 mm column using elution phases A (5% formic acid in water) and B (5% formic acid in acetonitrile). A step gradient was initiated with 1 min 95% solvent A, followed by 5 min 90% solvent A, and 75% solvent A and for 30 min. Finally, solvent A was gradually dropped to 5% over a period of 40 min and held for a further one minute. Detection via UV spectrometer was performed at 525 nm and quantified using the MRM method, with reference to an external standard (cyanidin 3.5-diglucoside).

### 2.8 Fermentation of single bacterial strains

Individual bacterial strains *Anaerobutyricum hallii* DSM 3353, *Anaerobutyricum soehngenii* L2-7, *Enterocloster bolteae* R7, *Flavonifractor plautii* DSM400, *Coprococcus* sp. L2-50 and *Fusicatenibacter saccharivorans* D5 BHI MAN 8 were subjected to further fermentation experiments to study the metabolism of catechin. The selected bacterial strains were cultivated anaerobically in triplicate at 37°C in 5 mL fermentation media (Duncan et al., 2016), containing either 3mg formulation extract (Cueva et al., 2013) or catechin standard (500µg/L). Additional anaerobic media controls containing either 3mg pre-digested formulation extract or catechin standard (500µg/mL) were analysed. Samples were taken at 0h and 24h via a syringe (300 µL) during CO_2_ gassing, and were analysed for flavonoid concentrations (catechin, epicatechin, and epigallocatechin) (Russell et al., 2011).

### 2.9 Statistical analysis

Statistical analysis and visualisation of data was performed in R version 4.1.1 (R Foundation for Statistical Computing, 2021). The package ggplot2 (Wickham, 2016) was used for generating figures. Normality was tested via Royston tests using the package MVN (Korkmaz et al., 2014). Phenolic metabolite composition from extract powders of the different phases in the *in vitro* digestion model were averages of biological duplicates and summarised phenolic content of three extraction stages per extract sample, expressed as the sum ± root mean square of phenolic content.

Overall gut microbiota composition was assessed via the parsimony and analysis of molecular variance (AMOVA), commands in mothur (Schloss et al., 2009). The Shannon bacterial diversity score was calculated in mothur, and statistical significance was evaluated via a two-way ANOVA and Tukey post-hoc test (factors were Shannon diversity scores, timepoints, and volunteer/control samples). ANOVA results were presented as F-test values (degrees of freedom) and p-value, according to Field et al. (2012). Statistical analysis of phenolic metabolite bioconversion in the fermentation experiment was performed by two-way ANOVA (covariates were phenolic metabolite concentration, factors were timepoints, or volunteer/control samples) by comparing timepoints (2h, 5h, 24h, 48h and 72h) to the baseline measurement, followed by Tukey post-hoc analysis. The SCFAs were statistically analysed via a linear mixed model with volunteer and fermentation flask as random effects, and extract/control conditions and timepoint as fixed effects (Kuznetsova et al., 2017; Searle et al., 1980), denominator degrees of freedom were obtained by Satterthwaite’s method. To assess the impact of presence or absence of the extract on the faecal bacterial composition during the incubation period (at 24h and 72h), Linear discriminant analysis Effect Size (LEfSe) (Segata et al., 2011) and Metastats (White et al., 2009) tools were employed in mothur (Schloss et al., 2009). LEfSe and Metastats results were corrected for multiple testing via the Benjamini-Hochberg method, with a false discovery rate of 0.05.

To analyse the relationship between faecal bacterial growth (increase in proportional abundance of bacterial Operational Taxonomic Units (OTUs) over time), and bioconversion of flavan-3-ols (decrease of (epi-)catechin content over time), a Pearson correlation was performed in Microsoft Excel for each individual donor. The p-values of the Pearson correlation were calculated in R using the corrplot function (Wei & Simko, 2024). The correlation data were analysed for catechin/epicatechin and bacterial OTU relationships, for each donor individually, to find negative correlations of R^2^<−0.7 (p <0.01), and were corrected for multiple testing via Benjamini-Hochberg testing, with a false discovery rate of 0.05.

The catechin levels in the presence of individual bacterial cultures (*Anaerobutyricum hallii* DSM3353, *Anaerobutyricum soehngenii* L2-7, *Coprococcus* sp. L2-50, *Enterocloster bolteae* R7, *Flavonifrac tor plautii* DSM4000, *Fusicatenibacter saccharivorans* D5 BHI MAN 8, and bacterial free control) were analysed via ANOVA, with factors being sample types (extract and catechin standard), timepoints (0h and 24h), and cultivation flasks (as a random effect). The epicatechin and epigallocatechin were only measured in the pre-digested formulation extract due to the lack of pure standards for comparison. The changes in epicatechin and epigallocatechin levels were analysed via the t.test function in R, comparing the individual flavonoid levels (epicatechin, or epigallocatechin levels) and timepoints (0h and 24h) for each individual bacterial culture (*Anaerobutyricum hallii* DSM3353, *Anaerobutyricum soehngenii* L2-7, *Coprococcus* sp. L2-50, *Entero closter bolteae* R7, *Flavonifractor plautii* DSM4000, *Fusicatenibacter saccharivorans* D5 BHI MAN 8, and bacterial free control).

## 3 Results

### 3.1 Composition of the commercial and formulation extracts

Grape seed extract contained the highest levels of monomeric flavan-3-ols (1760 ± 75ng / 100mg), whereas the bilberry extract had much higher levels of benzoic acids (496 ± 15ng / 100mg) compared to grape seed extract (219 ± 16ng / 100mg). Levels of conjugated (epi-)catechins (gallocatechin and epigallocatechin combined) were higher in the commercial bil berry extract compared to the grape seed extract (163.5 ± 36.7ng / 100mg versus 65.5 ± 19.9ng / 100mg). Flavonoids and relevant metabolites in the bilberry, grape seed and formulation extracts are presented in Figure 1 and in Supplementary Table 1.

**Figure 1.**
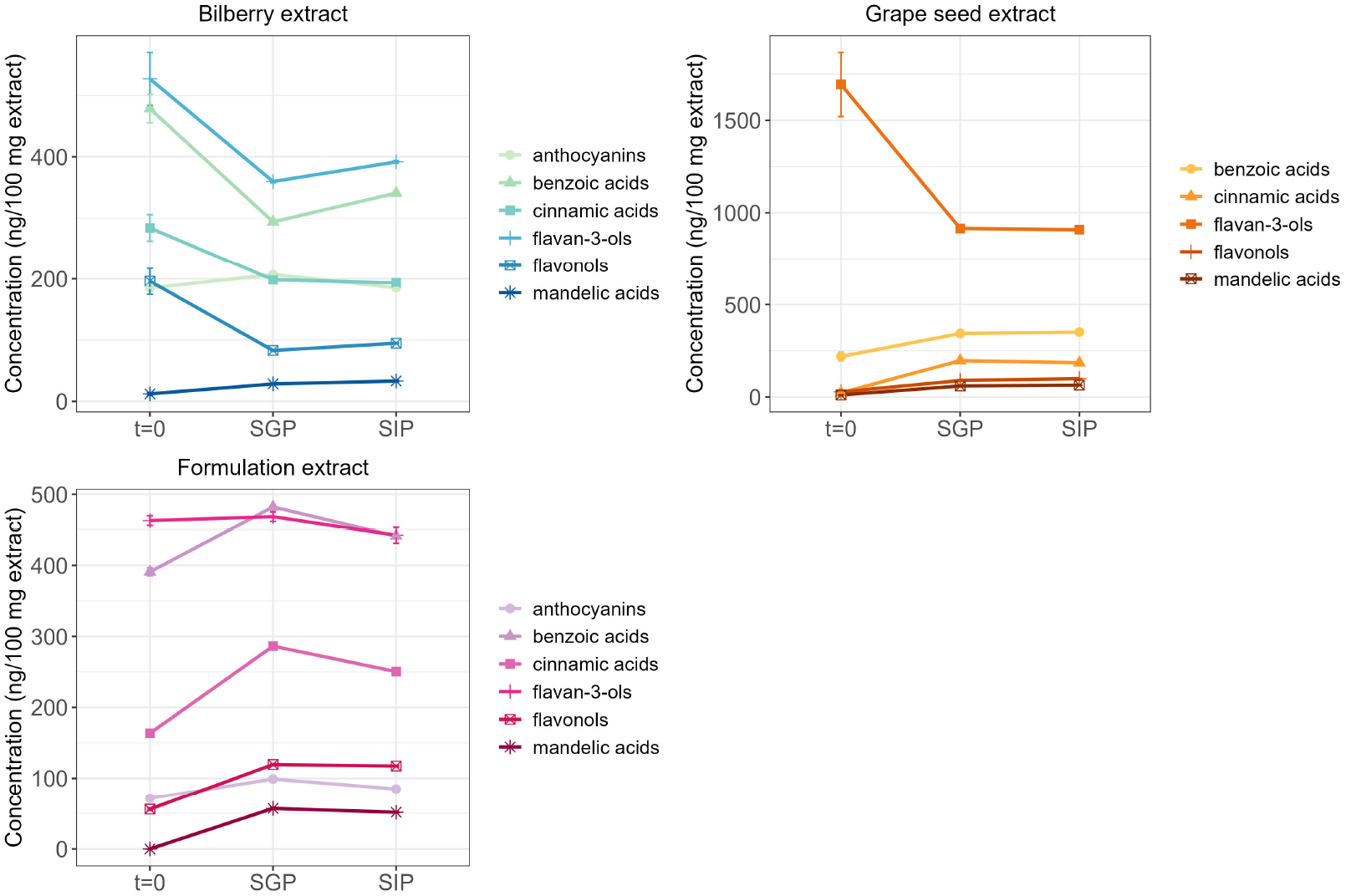
Phenolic metabolite content (of anthocyanins, benzoic acids, cinnamic acids, flavan-3-ols, flavonols and mandelic acids) in the bilberry, grape seed and formulation extracts (t = 0) and after gastric (SGP) and intestinal (SIP) *in vitro* digestion phases. Data are represented as sum ± root mean square. SGP – simulated gastric phase, SIP – simulated intestinal phase.

Whilst anthocyanins were detected in bilberry and formulation extract samples, grape seed ex tract did not contain any of the tested anthocyanins (cyanidin, delphinidin, malvidin, pelargoni din, peonidin, petunidin). The total anthocyanin content in bilberry extract sample was 18.5% (w/w), and in the formulation extract 7.2% (w/w). Levels of anthocyanins between the bilberry and the formulation extract were significantly different (F_(1,80)_ = 240.49, p<0.001) (Supplementary Table 2).

### 3.2 Flavonoid metabolism during *in vitro* simulated gastro-intestinal passage

Generally, levels of the flavanols catechin, epicatechin, epigallocatechin gallate and gallocatechin in grape seed and bilberry extract decreased after the combined salivary and gastric stage of digestion, with little further digestion during the intestinal phase (Figure 1, Supplementary Tables 3 and 4). The anthocyanin content remained stable during the different digestion stages in bilberry (sum of anthocyanins in digestive stages: initial concentration = 185µg/mg, gastric digestion = 207µg/mg, intestinal digestion = 186 µg/mg) and formulation extract (sum of anthocyanins in digestive stages: initial concentration = 71 µg/mg, gastric digestion = 99µg/mg, intestinal digestion = 85µg/mg).

We next investigated the potential impact of the gut microbiota on metabolism of flavonoids, by carrying out faecal incubations. As expected, each volunteer had a unique faecal microbial composition (parsimony analysis of Bray-Curtis dissimilarity dendrogram, p = 0.01 Figure 2), which was reflected in the overall dendrogram clustering into six discrete clusters of samples, each derived from individual donors.

**Figure 2.**
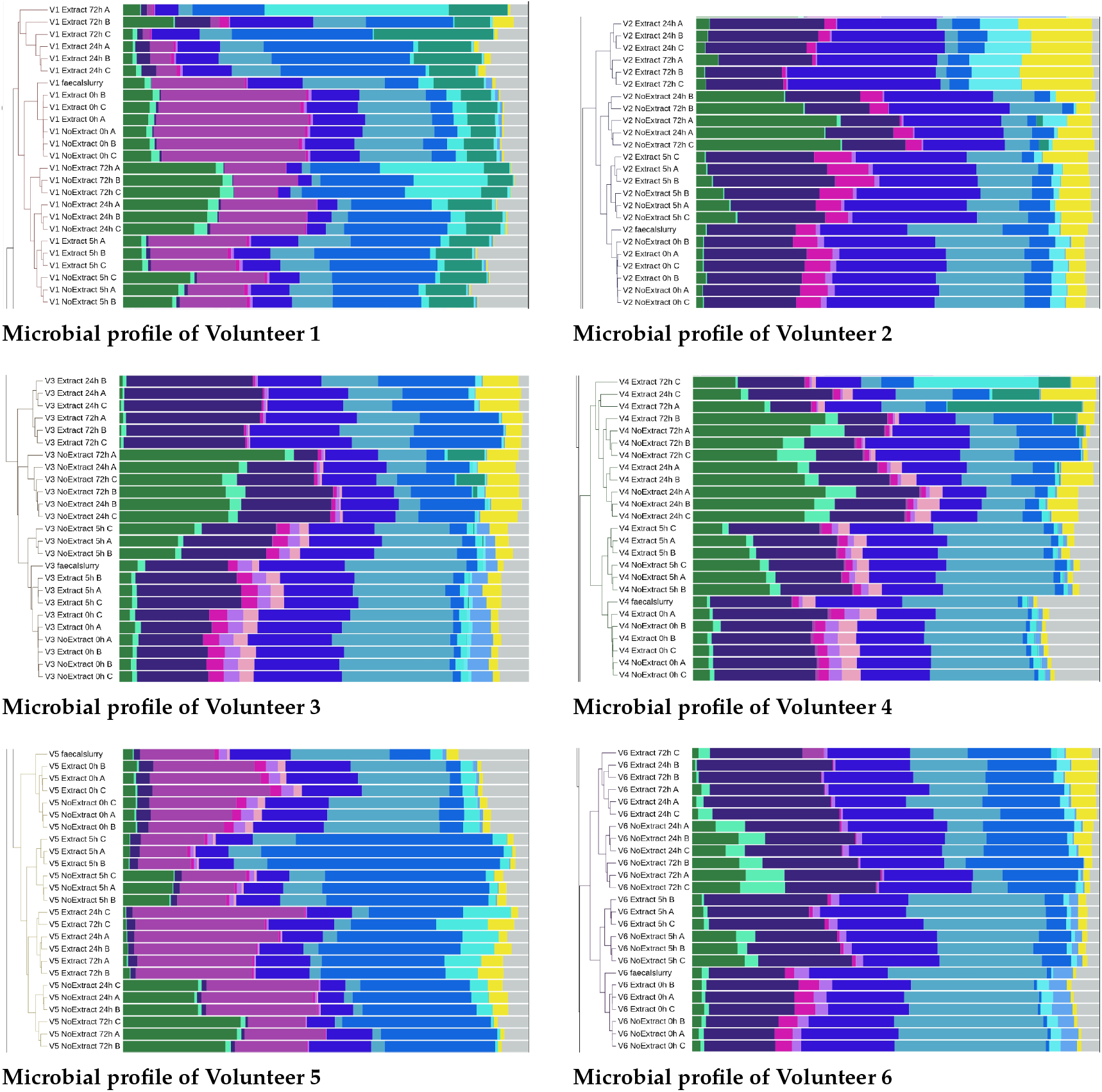
Overall shifts in faecal bacterial composition after incubation with the final pellet samples of the formulation extract, obtained from the *in vitro* digestion model, which were inoculated with faecal matter from six volunteers (V1 to V6). Samples were taken at baseline (t = 0h) and after 5h, 24h and 72h of incubation. Samples with the suffix A to C represent biological triplicates. The “faecal slurry” samples are the initial baseline faecal microbiota composition of the six donor samples. The proportional abundances of selected bacterial families are presented in the key to the right of the dendrogram and were coloured according to the phylum they belong to: Actinobacteria (in greens: 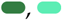), Bacteroides (in purples: 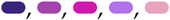), Firmicutes (in blues: 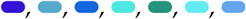), Proteobacteria (in yellow: 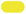), and others (in grey: 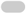). The complete iTOL figure is available in Supplementary Figure 11.

Significant changes in bacterial diversity, as assessed using the Shannon diversity index, were observed between the 0h and 72h time points for each faecal donor (Supplementary Figure 1). There were significant changes in the bacterial communities during fermentation with the formulation extract relative to the faecal control samples, suggesting that the presence of phenolic metabolites favoured the growth of certain bacteria. In the presence of the extract, donor samples contained higher proportional abundances of Firmicutes (in blues: 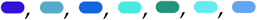); LEfSe Benjamini-Hochberg corrected p = 0.005, Metastats Benjamini-Hochberg corrected p = 0.008) and Proteobacteria (in yellow: 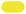); LEfSe Benjamini-Hochberg corrected p = 0.035, Metastats Benjamini-Hochberg corrected p = 0.020), while no-extract containing samples were proportionally enriched with Actinobacteria (in greens: 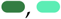); LEfSe Benjamini-Hochberg corrected p = 0.002, Metastats Benjamini-Hochberg corrected p = 0.002). Bacterial families significantly increased in proportional abundance in the presence of the extract were Ruminococcaceae 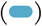 at 24h (Figure 2, second shade of blue; LEfSe Benjamini-Hochberg corrected p = 0.023, Metastats Benjamini-Hochberg corrected p = 0.038), and Sutterellaceae at 72h (Figure 2, yellow 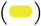; LEfSe Benjamini-Hochberg corrected p = 0.023, Metastats Benjamini-Hochberg corrected p = 0.017). Bifidobacteriaceae 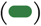 and Coriobacteriaceae 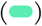 (LEfSe Benjamini-Hochberg corrected p = 0.002, Metastats Benjamini-Hochberg corrected p = 0.003; and LEfSe Benjamini-Hochberg corrected p = 0.006, Metastats Benjamini-Hochberg corrected p = 0.005 respectively) were found in significantly increased proportional abundance at 24h and 72h in the control samples.

Additionally, unique SCFA profiles from the faecal fermentations generated from the six volunteer samples, were measured. Propionate and acetate levels increased significantly from 0h to 24h, 0h to 48h and 0h to 72h in both extract and faecal control conditions across volunteers (F_(3,13)_ <0.0001, F_(3,93)_ = 0.002 respectively). Significantly higher levels of propionate were measured in the presence of the extract compared to the faecal control samples (F_(1,26)_ = 9.6, p = 0.005) at 48h to 72h (p = 0.002 and p = 0.001 respectively), suggesting that the phenolic metabolites present in the extract may favour the growth of certain propionate-producing bacteria (Supplementary Figure 2).

Next, the bioconversion of phenolic metabolites in the presence of the faecal microbiota was investigated. Levels of catechin, epicatechin and quercetin-3-glucoside decreased significantly during faecal fermentation, whilst levels in control samples remained stable, indicating a rapid metabolism of these flavonoids into derivative metabolites by faecal microbiota in all six donors (Figure 3). There was no gallocatechin detected throughout the fermentation experiment. Phenylpropionic, phenylacetic and phenyl lactic acid metabolism is depicted in Supplementary Figures 3 and 4. Cinnamic acids (Supplementary Figure 5) disappeared within 24h of inoculation in all donor samples indicating complete bioconversion by the faecal microbiota.

**Figure 3.**
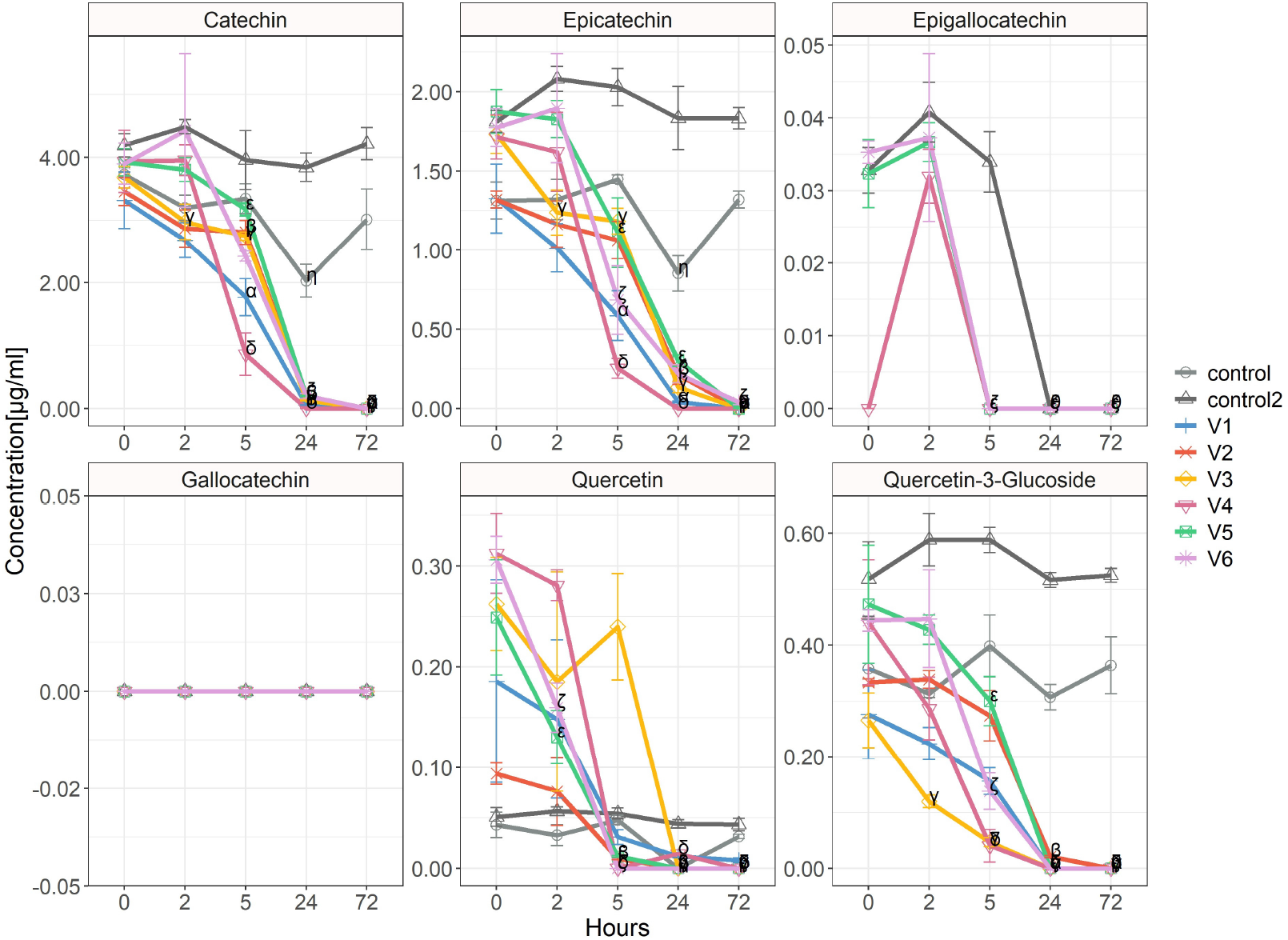
Bioconversion of flavan-3-ols and flavonoids by faecal microbiota present in the faecal samples of six healthy donors (V1 to V6), and two extract controls (containing only formulation extract and growth media but no faecal bacteria) at baseline (0h), and after 2h, 5h, 24h and 72h of inoculation. donor/sample specific p-values (<0.05): V1-α, V2-β, V3-γ, V4-δ, V5-ε, V6-ζ, control – η, control 2 – θ, compared to the baseline time point.

Within 24 h the flavan-3-ol and flavonoid content was depleted in the presence of faecal microbiota of all six donors, and the cultures were enriched with benzoic acids, phenylpropionic acids, phenyllactic acids, and phenylacetic acids. A comparable pattern of benzoic acid formation, especially that of gallic acid, protocatechuic acid, syringic acid and vanillic acid, was identified for each unique donor sample. The benzoic acid formation peaked between 5 h and 24 h for all donors (Figure 4), which suggests that the formation of these metabolites coincide with the breakdown of catechin and epicatechin.

**Figure 4.**
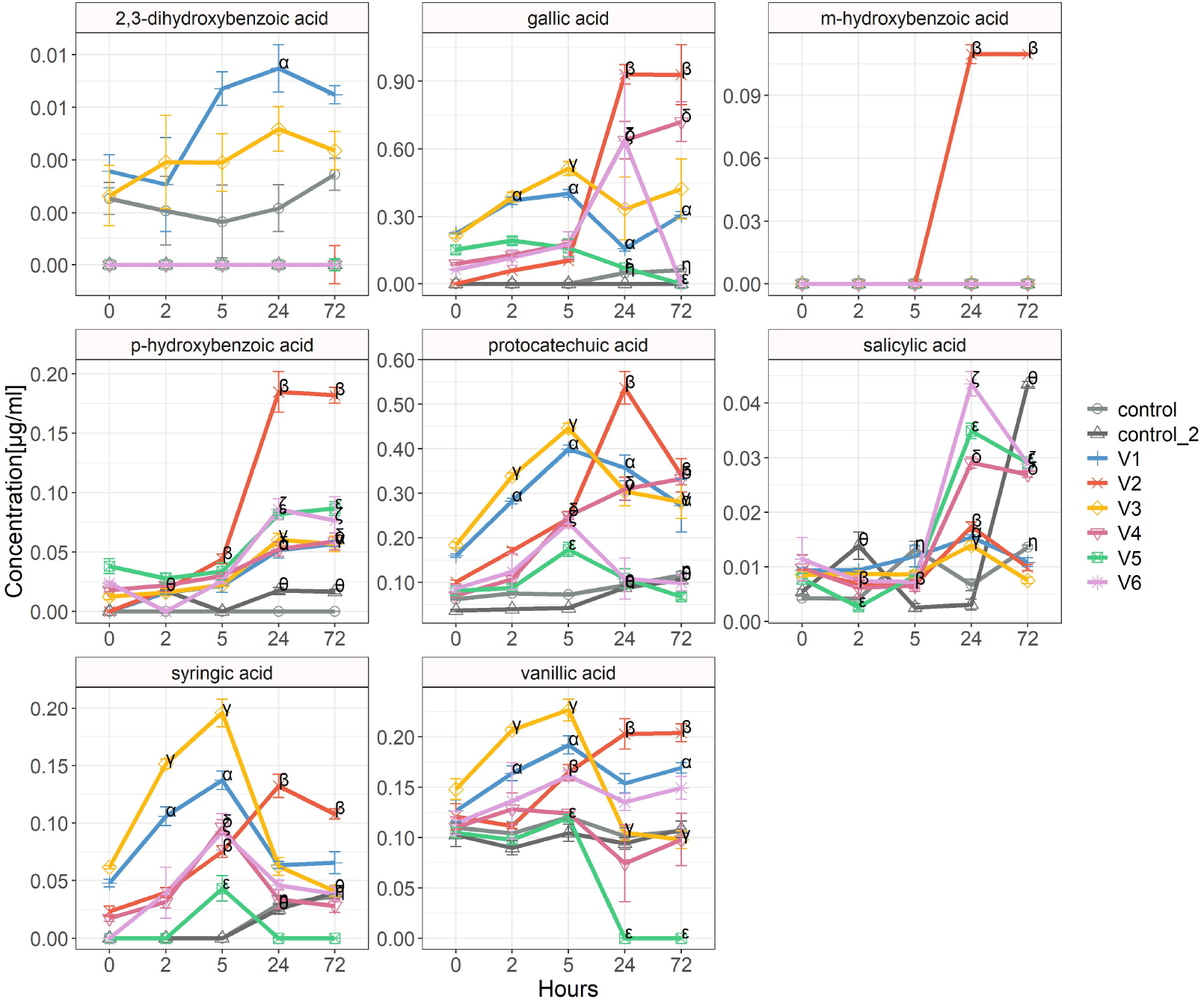
Benzoic acid formation by faecal microbiota present in the faecal samples of six healthy donors (V1-V6), and two extract controls (containing only formulation extract and growth media but no faecal bacteria) at baseline (0h), and after 2h, 5h, 24h and 72h of inoculation. donor/sample specific p-values (<0.05): V1-α, V2-β, V3-γ, V4-δ, V5-ε, V6-ζ, control – η, control 2 – θ, compared to the baseline time point.

OTUs related to the bacterium *Anaerobutyricum hallii*, which was identified in five out of six donors, and a previously uncultured species most closely related (96% similarity) to *Entero closter bolteae*, which was identified in four out of six donors, were both negatively correlated with the presence of (epi-)catechin indicating that both *Anaerobutyricum hallii* and *Enterocloster bolteae* may play a role in the bioconversion of (epi-)catechin to smaller phenolic metabolites. Correlation analysis identified 17 OTUs that were negatively correlated with the presence of catechin and epicatechin (R *≤*−0.7, p *≤*0.01) for donor V1. Most of these OTUs were found to belong to the Bacteroides genus (n=7) (Figure 5). Details of bacterial OTUs, negatively associated with catechin levels, for all other donors can be found in Supplementary Figures 6-10, and Supplementary Table 6.

**Figure 5.**
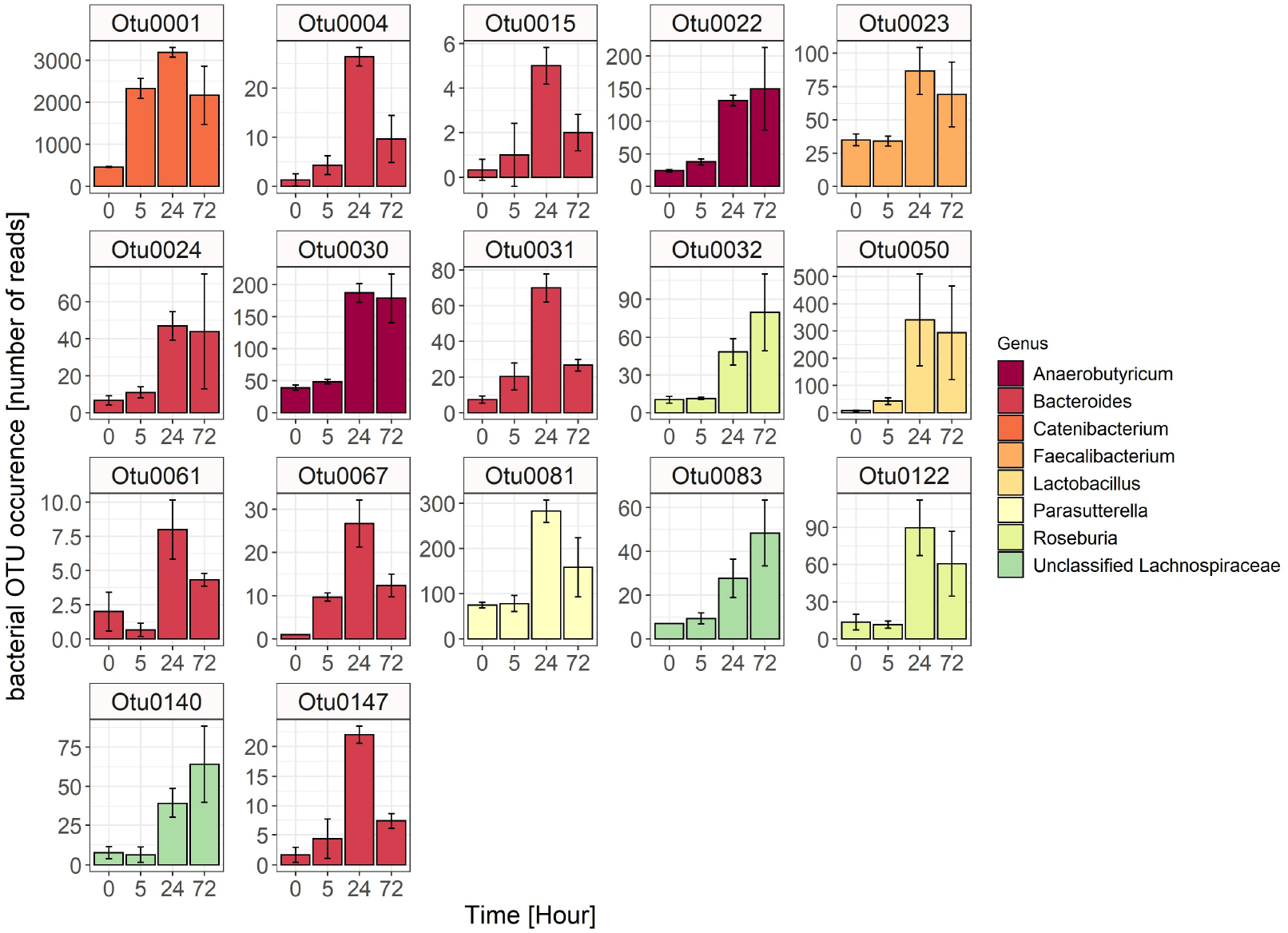
Bacterial OTUs from the faecal donor V1, which were negatively correlated with the presence of catechin and epicatechin from the formulation extract. Data are represented as mean ± standard deviation of biological triplicates.

To further investigate the hypothesis of single faecal bacterial involvement in catechin/epicatechin metabolism, fermentation experiments using single bacterial strains incubated with catechin standard and pre-digested formulation extract were performed. The bacterial stains were chosen based on the correlation analysis above, and on previous literature by Grohmann et al. (2023) and Kutschera et al. (2011). Catechin standard levels did not significantly change in any of the single bacterial cultures at 24h. However, catechin levels in the formulation extract increased significantly in the presence of *Anaerobutyricum hallii DSM3353* (F_(1,7)_ = 17.4, p = 0.004) at the 24h timepoint, while concentrations did not significantly change in the presence of other single bacterial cultures (Figure 6). Furthermore, no significant changes in epicatechin and epigallocatechin levels were detected in the formulation extract samples, in the presence of single bacterial cultures (Supplementary Table 7).

**Figure 6.**
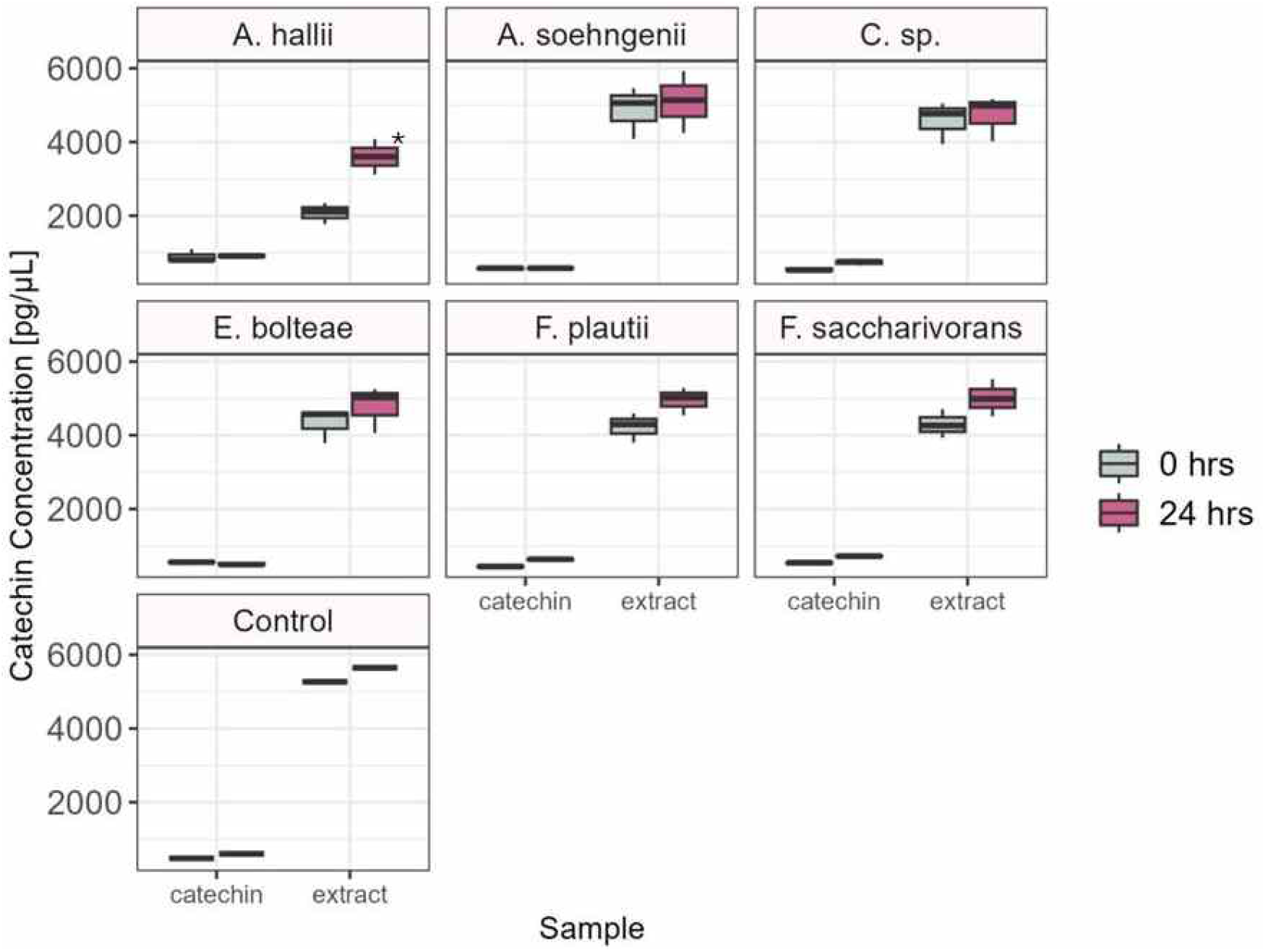
Single bacterial strains (*Anaerobutyricum hallii* DSM3353, *Anaerobutyricum soehngenii* L2-7, *Coprococcus* sp. L2-50, *Enterocloster bolteae* R7, *Flavonifractor plautii* DSM4000, *Fusicatenibacter saccharivorans* D5 BHI MAN 8 in triplicate, and bacterial free control) were inoculated with pre-digested formulation extract (3mg/5mL) or catechin standard (500 µg/mL) and cultivated anaerobically. The media control only contained pre-digested formulation extract (3mg/5mL) or catechin standard (500 µg/mL) and no bacteria. *∗*-significant difference between 0h and 24h, p<0.05.

## 4 Discussion

### 4.1 Phenolic composition of bilberry extract, grape seed extract and formulation extract

This work sought to study changes to the phenolic profile of a bilberry, grape seed, and a formulation extract as they passed along the different phases of an *in vitro* simulated digestion model, followed by a faecal fermentation. At the simulated salivary and gastric phases, levels of flavan-3-ol decreased in all three extracts, while the content of smaller metabolites such as cinnamic acids, benzoic acids and mandelic acids increased.

The model was extended further to include the use of faecal samples from six healthy donors to study the effects of colonic microbial metabolism on the formulation extract fraction obtained from the intestinal phase. Unique compositions of faecal microbiota from six healthy donors were capable of metabolising flavan-3-ols in the formulation extract digesta within 24h, result ing in higher concentrations of benzoic acids, phenylpropionic acids, phenylacetic acids, and propionate. Flavan-3-ol metabolism appeared to be linked to an increased relative abundance of *Anaerobutyricum hallii* and *Enterocloster bolteae* across five faecal donors. However, fermentation using pure bacterial cultures with the formulation extract and a catechin standard found that they were incapable of monomeric catechin metabolism.

The analysis of the phenolic content of grape seed and bilberry extract indicated that they are rich in flavan-3-ols and benzoic acids. Overall, phenolic content increased in the formulation extract after the gastric digestion phase and remained stable in the intestinal digestion phase, while the phenolic content of bilberry and grape seed extracts decreased after the gastric phase and remained stable in the intestinal phase. In bilberry and grape seed extracts, a decrease of monomeric (epi-)catechins was detected comparing the initial extract composition to the gastric phase, but no further decrease in levels of monomeric (epi-)catechins was observed between the gastric and intestinal phase (at pH neutral conditions). In the absence of bacteria, the decay of polymeric flavan-3-ols is expected to occur in acidic environments, whereas the monomeric flavan-3-ols and anthocyanins decay at pH 7 to 8.5 (Cheynier et al., 2012; Williamson et al., 2018). Of course, variances can be expected when comparing phenolic metabolite content between static, semi-static and dynamic models. While dynamic models such as those employed by Dupont et al. (2019), were able to replicate physiological conditions more closely, static models tend to have the advantage of being far more convenient to control and generate samples consistently. For example, Barik et al. (2020) who employed a static model demonstrated a reduction of phenolic compound levels by 60% and 96% in blackcurrant and greencurrant extract after the intestinal digestion phase, employing the same *in vitro* digestion model as described in this work. Similarly, an *in vitro* digestion of grape seed extract performed by Wang et al. (2013) showed an initial increase in catechin, epicatechin and gallic acid levels after the gastric phase that also significantly dropped after the pancreatic/intestinal phase. Overall differences in study outcomes though are likely a function of the varying feed material (grape seed, bilberry, blackcurrant, greencurrant etc), or even the natural variation in the digestive porcine enzymes employed in the model, or the solubility of the extracts in the *in vitro* digestion vessels impacting the exposure to pH and enzyme conditions.

In the work described herein, the difference in the anthocyanin levels in bilberry and formulation extract across the various stages of the simulated digestion model were marginal. This contrasts with the decrease in anthocyanin levels observed by Barik et al. (2020), Corrêa et al. (2017), and Lingua et al. (2018). The lack of change in the monomeric anthocyanin content could be due to the degradation of the multimeric forms of the compound, which were not measured in this experiment, that kept replenishing anthocyanin level resulting in a net zero change. Moreover, a closely pH-monitored and anaerobic environment might have been more suitable for the analysis of anthocyanin metabolism. For example, Iyer et al. (2023) described a controlled semi-static model incorporating the use of anaerobic condition during the gastric and intestinal phase to replicate the physiological conditions, in addition to the use of cell culture models as well as faecal fermentation to provide a closer approximation to physiological conditions.

The faecal samples of six healthy donors harboured unique sets of intestinal microbiota, which nonetheless led to the complete catabolism of flavonoids to phenolic acid metabolites, such as phenylpropionic, phenylacetic and benzoic acids, within 24h in every case. Interestingly, a study by Xu et al. (2024) found that the metabolism of catechin and epicatechin from grape seed extract can be enhanced in the presence of probiotic bacteria. Similarly, live faecal bacteria in an anaerobic and pH-controlled fermentation system metabolised anthocyanins from a bilberry extract within eight hours, while anthocyanins decayed slower in a fermentation system without live bacteria (Shehata et al., 2023). Although anthocyanin metabolism was not measured in the faecal fermentation experiment described here, it is plausible that anthocyanins in the digested formulation extract samples (estimated anthocyanin content in the formulation extract of ~7.2% w/w, Section 3.1) will have contributed to the formation of phenolic acids. In previous fermentation experiments, the anthocyanin-glucoside content was depleted within 4h to 8h in the presence of gut microbiota, while anthocyanins with more complex sugar moieties, such as arabinose, were still detectable after 24h (Wu et al., 2020).

We hypothesised that the faecal bacteria whose abundance negatively correlated with the concentration of catechin and epicatechin, and positively correlated with the concentration of phenolic acid metabolites, could plausibly be involved in catechin and epicatechin metabolism. The exploratory correlative 16S rRNA gene analysis suggested that two microbial species, namely, *Anaerobutyricum hallii* and *Enterocloster bolteae* would be worthy of further investigation (Figure 5, Supplementary Figures 6-10). However, previous studies on phenolic metabolite conversion have not identified *Anaerobutyricum hallii* and *Enterocloster bolteae* as potential candidates for catechin and epicatechin metabolism, although this may be complicated by the fact that both species have recently been taxonomically re-classified (Haas & Blanchard, 2020; Shetty et al., 2018). Conversely, while other studies have identified that *Eggerthella lenta* in conjunction with *Flavonifractor plautii* can metabolise (epi-)catechins to 5-(3,4-dihydroxyphenyl)-c-valerolactone and 4-hydroxy-5-(3,4-dihydroxyphenyl)valeric acid (Kutschera et al., 2011), neither of these two species were detected in this *in vitro* study, possibly because they were not present in significant abundance in any of the faecal donor samples. Importantly, our follow-up fermentation experiments with single bacterial strains with catechin standard and pre-digested formulation extract indicated that none of the faecal bacterial strains of interest (*Anaerobutyricum hallii* DSM3353, *Anaerobutyricum soehngenii* L2-7, *Coprococcus* sp. L2-50, *Enterocloster bolteae* R7, *Flavonifractor plautii* DSM4000, *Fusicatenibacter saccharivorans* D5 BHI MAN 8) were capable of metabolising monomeric catechin. However, the significant increase in catechin levels in the presence of *Anaerobutyricum hallii* could indicate the ability to metabolise polymeric flavonoids. A study by Di Pede et al. (2022) was able to show that polymeric flavonoids are being metabolised by faecal microbiota in an *in vitro* fermentation, although specific species were not identified. Although we were only testing the association of these faecal bacteria with (epi-)catechin level reduction, they could have been involved in the formation of benzoic acids, which positively correlated with the bacterial presence and negatively correlated with the concentration of (epi-)catechins. Further *in vitro* studies, single strain fermentation experiments, but also *in vivo* and human intervention studies are needed to identify faecal bacteria capable of phenolic metabolism.

### 4.2 Strengths and limitations to the study

The strengths of this work are the detailed analysis of the *in vitro* metabolism of grape seed, bilberry and formulation extracts in a simulated digestion, followed by faecal fermentation and single bacterial cultures. This work supported the current literature on phenolic metabolism and highlighted the importance of faecal microbiota to completely metabolise flavan-3-ols. The rigorous faecal fermentation setup and controls enabled the monitoring of metabolism of phenolic compounds, but also indicated which phenolic metabolites were likely remnants of the individual donor’s diet.

The *in vitro* digestion and faecal fermentation models employed in this work have certain limitations. Firstly, polymeric molecules could not be measured, and consequently, mass balance to track the increase in monomeric catechin/epicatechin concentrations was not possible. Consequently, the polymeric (epi-)catechins could have been metabolised into monomers in the initial stages of the faecal fermentation experiment. The majority of flavonoids in fruits are present in polymeric form, comprising up to 74% of a fruits total flavonoid content (Kelm et al., 2004). We also found that the acidic and basic environments during the phenolic extraction procedure, as well as the *in vitro* digestion processes, were not sufficient to metabolise monomeric flavan-3-ols and anthocyanins into smaller phenolic acids. As indicated above, this might be explained by the static *in vitro* model employed. A further limitation of our *in vitro* digestion and faecal fermentation is the unnatural accumulation of phenolic metabolites and SCFAs, that may otherwise be transported across the intestinal lumen *in vivo*. Consequently, their accumulation might have had an impact on the pH environment and thereby on the microbial community, and the phenolic metabolite profile.

## 5 Conclusion and future work

In this study we found that *in vitro* digestion of a bilberry, grape seed or a combined formulation extract decreased the content of catechin/epicatechin and increased the content of smaller phenolic metabolites such as benzoic acids, cinnamic acids and mandelic acids, whilst the anthocyanin content remained relatively stable. Subsequent incubation studies, where individual faecal microbiota from six donors were inoculated with the solid fraction end-product of the formulated extract after the *in vitro* digestion, saw the formation of unique sets of phenolic metabolites from catechin/epicatechin metabolism. We found that the formation of the smaller metabolites (phenylacetic acids, phenylpropionic acids, phenyllactic acid and benzoic acids) was correlated with the presence of certain bacterial taxa such as *Anaerobutyricum*, and *Enterocloster*. This work showed that the single bacterial strains *Anaerobutyricum hallii* DSM3353, *Anaerobutyricum soehngenii* L2-7, *Coprococcus* sp. L2-50, *Enterocloster bolteae* R7, *Flavonifractor plautii* DSM4000, and *Fusicatenibacter saccharivorans* D5 BHI MAN 8 are not capable of metabolising monomeric (epi-)catechins to smaller phenolic compounds. This work adds further proof that the baseline microbiota composition, and its metabolic activities, could impact the bioavailability of the phenolic metabolites, and therefore govern the inter-individual response to a bilberry-grape seed intervention.

## Acknowledgements

The authors wish to acknowledge the faecal sample donors for participating in the study, and Sylvia Duncan, Lorraine Scobbie, Gary Duncan, and Donna Henderson for technical assistance. PAK and MP gratefully acknowledge the support of the Biotechnology and Biological Sciences Re search Council (BBSRC) who funded this research via the BBSRC Institute Strategic Programmes ‘Food Microbiome and Health’ BB/X011054/1 and its constituent projects BBS/E/QU/230001A and BBS/E/QU/230001B and ‘Food Innovation and Health’ BB/R012512/1 and its constituent projects BBS/E/F/000PR10345 and BBS/E/F/000PR10346.

