## Supplementary_file for "Assessment of inter-individual variation in metabolism of flavonoids from bilberry and grape seed extracts using an *in vitro* digestion and faecal fermentation model"

Supplementary Table 1: Content of flavonoids and metabolites in the commercial bilberry, grape seed, and combination extracts.

|  | Bilberry extract (Mirtoselect®) | Grape seed extract (Enovita®) | Formulation extract* |
| --- | --- | --- | --- |
|  | (*ng/100 mg extract*) | | |
| flavonoids/coumarins | **953.8 ± 20.7** | **1760.0 ± 74.6** | **603.1 ± 3.2** |
| catechin | 168.6 ± 56.6 | 1074.6 ± 325.1 | 300.5 ± 12.1 |
| epicatechin | 195.0 ± 69.0 | 554.2 ± 209.6 | 132.6 ± 9.0 |
| gallocatechin | 65.5 ± 15.5 | 43.4 ± 19.1 | 10.0 ± 1.0 |
| epigallocatechin | 97.9 ± 33.3 | 1.4 ± 0.7 | 17.6 ± 2.0 |
| epigallocatechin gallate | 0.0 ± 0.0 | 20.7 ± 5.5 | 2.3 ± 0.3 |
| quercetin | 190.9 ± 47.6 | 21.5 ± 7.0 | 24.5 ± 3.1 |
| isoliquiritigenin | 0.2 ± 0.1 | 0.0 ± 0.0 | 0.0 ± 0.0 |
| phloretin | 4.7 ± 1.2 | 0.3 ± 0.1 | 0.3 ± 0.1 |
| naringenin | 0.1 ± 0.0 | 1.5 ± 0.7 | 2.4 ± 0.4 |
| naringin | 0.1 ± 0.0 | 0.0 ± 0.0 | 0.0 ± 0.0 |
| hesperitin | 1.2 ± 0.3 | 0.0 ± 0.0 | 0.0 ± 0.0 |
| kaempferol | 72.9 ± 15.0 | 2.3 ± 1.0 | 1.7 ± 0.4 |
| myricetin | 80.1 ± 20.0 | 0.9 ± 0.4 | 51 ± 6.5 |
| quercetin-3-glucoside | 5.2 ± 1.8 | 6.0 ± 2.6 | 31.4 ± 1.9 |
| taxifolin | 1.4 ± 0.6 | 28.6 ± 10.6 | 5.7 ± 0.9 |
| genstein | 0.0 ± 0.0 | 0.0 ± 0.0 | 0.0 ± 0.0 |
| scopoletin | 0.1 ± 0.0 | 0.0 ± 0.0 | 0.8 ± 0.0 |
| umbelliferone | 0.1 ± 0.1 | 0.0 ± 0.0 | 0.0 ± 0.0 |
| 7,8-dihydroxy-6-methyl coumarin | 36.4 ± 15.1 | 0.0 ± 0.0 | 0.1 ± 0.0 |
| quercitrin | 0.3 ± 0.2 | 0.7 ± 0.3 | 10.2 ± 1.3 |
| biochanin a | 0.0 ± 0.0 | 0.0 ± 0.0 | 0.0 ± 0.0 |
| didymin | 8.8 ± 3.9 | 0.0 ± 0.0 | 0.0 ± 0.0 |
| phloridzin | 0.0 ± 0.0 | 1.8 ± 0.8 | 3.2 ± 0.4 |
| daidzein | 0.5 ± 0.1 | 0.0 ± 0.0 | 0.0 ± 0.0 |
| luteolin | 4.5 ± 0.7 | 0.5 ± 0.2 | 0.8 ± 0.1 |
| isorhamnetin | 0.0 ± 0.0 | 0.9 ± 0.4 | 1.9 ± 0.4 |
| formononetin | 0.0 ± 0.0 | 0.0 ± 0.0 | 0.0 ± 0.0 |
| apigenin | 0.1 ± 0.0 | 0.1 ± 0.0 | 0.1 ± 0.0 |
| gossypin | 19.1 ± 4.5 | 0.5 ± 0.2 | 5.8 ± 0.6 |
| rutin | 0.0 ± 0.0 | 0.0 ± 0.0 | 0.0 ± 0.0 |
| benzoic acids | **495.6 ± 15.0** | **219.4 ± 15.6** | **396.3 ± 3.9** |
| salicylic acid | 15.0 ± 3.0 | 0.2 ± 0.1 | 5.0 ± 0.6 |
| m-hydroxybenzoic acid | 2.8 ± 1.1 | 0.0 ± 0.0 | 0.8 ± 0.1 |
| p-hydroxybenzoic acid | 20.6 ± 5.4 | 3.1 ± 0.6 | 7.5 ± 0.5 |
| 2,3-dihydroxybenzoic acid | 1.8 ± 0.8 | 0.0 ± 0.0 | 0.4 ± 0.0 |
| 2,4-dihydroxybenzoic acid | 0.0 ± 0.0 | 0.1 ± 0.0 | 0.0 ± 0.0 |
| 2,5-dihydroxybenzoic acid | 15.2 ± 6.7 | 0.0 ± 0.0 | 4.3 ± 0.2 |
| 2,6-dihydroxybenzoic acid | 0.0 ± 0.0 | 0.0 ± 0.0 | 0.7 ± 0.1 |
| protocatechuic acid | 188.2 ± 32.6 | 45.0 ± 14.6 | 86.6 ± 7.6 |
| p-anisic acid | 16.3 ± 3.1 | 0.9 ± 0.2 | 2.6 ± 0.2 |
| gallic acid | 74.0 ± 17.3 | 166.8 ± 52.0 | 192.2 ± 8.9 |
| vanillic acid | 72.6 ± 15.2 | 1.9 ± 0.3 | 34.1 ± 1.9 |
| syringic acid | 89.2 ± 31.8 | 1.3 ± 0.3 | 62.0 ± 4.9 |
| benzaldehydes | **28.5 ± 2.0** | **93.3 ± 9.5** | **36.7 ± 0.7** |
| p-hydroxybenzaldehyde | 1.0 ± 0.1 | 1.1 ± 0.3 | 0.5 ± 0.1 |
| protocatachaldehyde | 14.3 ± 3.4 | 85.2 ± 23.2 | 29.7 ± 1.5 |
| 3,4,5-trihydroxybenzaldehyde | 11.4 ± 3.7 | 2.6 ± 1.2 | 4.2 ± 1.0 |
| vanillin | 0.6 ± 0.0 | 4.3 ± 1.4 | 2.0 ± 0.3 |
| syringin | 1.2 ± 0.2 | 0.1 ± 0.0 | 0.2 ± 0.0 |
| 3-methoxybenzaldehyde | 0.0 ± 0.0 | 0.1 ± 0.0 | 0.0 ± 0.0 |
| 20hbaldc | 0.0 ± 0.0 | 0.0 ± 0.0 | 0.1 ± 0.0 |
| cinnamic acids | **306.5 ± 15.6** | **22.8 ± 1.3** | **169.6 ± 2.0** |
| cinnamic acid | 2.2 ± 0.8 | 1.8 ± 0.5 | 2.1 ± 0.3 |
| o-coumaric acid | 6.2 ± 2.7 | 0.0 ± 0.0 | 1.3 ± 0.0 |
| p-coumaric acid | 120.0 ± 38.8 | 4.3 ± 1.1 | 88.2 ± 4.4 |
| caffeic acid | 119.4 ± 28.4 | 0.9 ± 0.2 | 63.6 ± 4.6 |
| ferulic acid | 32.8 ± 5.6 | 15.4 ± 3.7 | 7.1 ± 0.5 |
| sinapic acid | 3.4 ± 0.7 | 0.2 ± 0.0 | 1.3 ± 0.1 |
| 3-methoxycinnamic acid | 0.0 ± 0.0 | 0.0 ± 0.0 | 0.0 ± 0.0 |
| 4-methoxycinnamic acid | 21.6 ± 9.0 | 0.2 ± 0.1 | 5.7 ± 0.4 |
| 3,4-dimethoxycinnamic acid | 0.8 ± 0.3 | 0.0 ± 0.0 | 0.3 ± 0.0 |
| 3,4,5-trimethoxycinnamic acid | 0.1 ± 0.0 | 0.1 ± 0.0 | 0.1 ± 0.0 |
| phenylpropionic acids | **8.1 ± 0.7** | **1.3 ± 0.3** | **3.0 ± 0.1** |
| phenylpropionic acid | 0.0 ± 0.0 | 0.5 ± 0.3 | 0.4 ± 0.1 |
| 2-hydroxyphenylpropionic acid | 0.1 ± 0.1 | 0.0 ± 0.0 | 0.0 ± 0.0 |
| 3-hydroxyphenylpropionic acid | 0.6 ± 0.3 | 0.0 ± 0.0 | 0.3 ± 0.0 |
| 4-hydroxyphenylpropionic acid | 2.6 ± 0.6 | 0.0 ± 0.0 | 0.6 ± 0.0 |
| 3,4-dihydroxyphenylpropionic acid | 2.1 ± 0.7 | 0.0 ± 0.0 | 0.0 ± 0.0 |
| 4-hydroxy-3-methoxyphenylpropionic acid | 2.7 ± 1.3 | 0.8 ± 0.4 | 1.7 ± 0.1 |
| benzenes | **11.2 ± 2.8** | **5.0 ± 1.7** | **4.4 ± 0.3** |
| 1,2-hydroxybenzene | 10.5 ± 3.9 | 5.0 ± 2.4 | 4.4 ± 0.4 |
| 1,2,3-trihydroxybenzene | 0.6 ± 0.3 | 0.0 ± 0.0 | 0.0 ± 0.0 |
| acetophenones | **0.7 ± 0.1** | **0.3 ± 0.1** | **0.4 ± 0.0** |
| 4-hydroxyacetophenone | 0.4 ± 0.1 | 0.1 ± 0.0 | 0.1 ± 0.0 |
| 4-hydroxy-3-methoxyacetophenone | 0.3 ± 0.1 | 0.3 ± 0.1 | 0.4 ± 0.0 |
| 4-hydroxy-3,5-dimethoxyacetophenone | 0.0 ± 0.0 | 0.0 ± 0.0 | 0.0 ± 0.0 |
| phenylacetic acids | **12.5 ± 2.2** | **42.6 ± 10.8** | **13.5 ± 2.3** |
| phenylacetic acid | 0.5 ± 0.1 | 0.2 ± 0.1 | 0.4 ± 0.1 |
| 4-hydroxyphenylacetic acid | 9.1 ± 3.6 | 1.3 ± 0.6 | 4.7 ± 0.4 |
| 3,4-dihydroxyphenylacetic acid | 2.9 ± 1.1 | 41.1 ± 18.6 | 8.3 ± 4.0 |
| mandelic acids | **12.3 ± 2.6** | **10.8 ± 1.9** | **0.0 ± 0.0** |
| 3-hydroxymandelic acid | 10.2 ± 4.4 | 1.1 ± 0.3 | 0.0 ± 0.0 |
| 4-hydroxymandelic acid | 0.0 ± 0.0 | 3.3 ± 1.5 | 0.0 ± 0.0 |
| 3,4-dihydroxymandelic acid | 2.1 ± 0.8 | 6.4 ± 3.0 | 0.0 ± 0.0 |
| phenyllactic acids | | | |
| 4-hydroxyphenyllactic acid | 13.0 ± 3.9 | 0.5 ± 0.2 | 4.3 ± 0.3 |
| phenolics others | **280.6 ± 59.3** | **1.1 ± 0.2** | **235.7 ± 18.4** |
| chlorogenic acid | 250.5 ± 118.0 | 0.0 ± 0.0 | 224.3 ± 36.7 |
| ethylferulate | 0.1 ± 0.1 | 0.5 ± 0.2 | 0.4 ± 0.1 |
| tyrosol | 5.0 ± 1.8 | 0.6 ± 0.3 | 2.6 ± 0.2 |
| hydroxytyrosol | 25.0 ± 11.7 | 0.0 ± 0.0 | 8.3 ± 0.6 |
| phenolic dimers | | | |
| resveratrol | 0.1 ± 0.0 | 3.1 ± 1.4 | 2.3 ± 0.4 |
| indoles | **71.9 ± 15.5** | **20.6 ± 3.9** | **0.2 ± 0.0** |
| indole-3-acetic acid | 0.1 ± 0.0 | 0.1 ± 0.1 | 0.2 ± 0.0 |
| indole-3-carboxylic acid | 71.8 ± 21.9 | 20.5 ± 5.6 | 0.0 ± 0.0 |

Top row in bold indicates the sum ± root mean square of the phenolic metabolite group within bilberry, grape seed and formulation extract. Individual phenolic metabolite data are presented as means ± standard deviation of the biological duplicates and sum of the phenolic metabolite content from all three extraction stages, in 100 mg of the original extract sample, and were rounded up. *54% Mirtoselect® + 46% Enovita®.

Supplementary Table 2: The anthocyanin ratios in bilberry and formulation extract samples

| Anthocyanin | Content (%) |
| --- | --- |
| cyanidin -3-o-galactoside | 9.3 |
| cyanidin -3-o-glucoside | 10.7 |
| cyanidin -3-o-pentoside | 9.1 |
| delphinidin -3-o-galactoside | 8.7 |
| delphinidin -3-o-glucoside | 9.1 |
| delphinidin -3-o-pentoside | 7.3 |
| malvidin -3-o-galactoside | 4.3 |
| malvidin -3-o-glucoside | 9.5 |
| malvidin -3-o-pentoside | 3.7 |
| pelargonidin -3-o-galactoside | 0.1 |
| pelargonidin -3-o-glucoside | 0.2 |
| pelargonidin -3-o-pentoside | BDL |
| peonidin -3-o-galactoside | 3.2 |
| peonidin -3-o-glucoside | 6.9 |
| peonidin -3-o-pentoside | 1.6 |
| petunidin -3-o-galactoside | 4.5 |
| petunidin -3-o-glucoside | 7.8 |
| petunidin -3-o-pentoside | 3.9 |

Anthocyanin content in a 100 mg extract sample, represented as percentage content (w/w), of both bilberry and formulation extract. BDL= below detection level

Supplementary Table 3: Flavonoid and metabolite content of bilberry extract

| BILBERRY |  |  |  |  |  |  |  |
| --- | --- | --- | --- | --- | --- | --- | --- |
| ng/100 mg extract | **raw extract** | **gastric phase** | **gastric phase** | **gastric phase** | **intestinal phase** | **intestinal phase** | **intestinal phase** |
| **Flavanoids/Coumarins** |  | **Pellet** | **supernatant** | **sum** | **pellet** | **Supernatant** | **sum** |
| Catechin | 168.6 ± 56.6 | 43.1 ± 1.0 | 59.1 ± 1.7 | 102.3 ± 1.9 | 48.2 ± 3.4 | 62.4 ± 0.8 | 110.6 ± 3.5 |
| Epicatechin | 195.0 ± 69.0 | 53.2 ± 0.8 | 84.1 ± 2.2 | 137.3 ± 2.3 | 65.1 ± 2.8 | 89.5 ± 0.7 | 154.6 ± 2.9 |
| Gallocatechin | 65.5 ± 15.5 | 11.7 ± 0.2 | 31.7 ± 0.9 | 43.4 ± 0.9 | 16.2 ± 1.0 | 29.3 ± 0.3 | 45.6 ± 1.1 |
| Epigallocatechin | 97.9 ± 33.3 | 20.9 ± 0.5 | 55.8 ± 2.3 | 76.8 ± 2.3 | 27.2 ± 1.4 | 53.9 ± 1.1 | 81.1 ± 1.8 |
| Epigallocatechin Gallate | 0.0 ± 0.0 | 0.0 ± 0.0 | 0.0 ± 0.0 | 0.0 ± 0.0 | 0.0 ± 0.0 | 0.0 ± 0.0 | 0.0 ± 0.0 |
| Quercetin | 190.9 ± 47.6 | 11.1 ± 0.4 | 21.2 ± 0.2 | 32.3 ± 0.4 | 17.5 ± 0.6 | 22.1 ± 0.7 | 39.6 ± 1.0 |
| Isoliquiritigenin | 0.2 ± 0.1 | 0.0 ± 0.0 | 0.0 ± 0.0 | 0.1 ± 0.0 | 0.0 ± 0.0 | 0.0 ± 0.0 | 0.1 ± 0.0 |
| Phloretin | 4.7 ± 1.2 | 0.1 ± 0.0 | 0.1 ± 0.0 | 0.3 ± 0.0 | 0.2 ± 0.0 | 0.1 ± 0.0 | 0.3 ± 0.0 |
| Naringenin | 0.1 ± 0.0 | 4.4 ± 0.2 | 3.5 ± 0.1 | 7.9 ± 0.2 | 4.6 ± 0.2 | 2.9 ± 0.0 | 7.5 ± 0.2 |
| Naringin | 0.1 ± 0.0 | 0.1 ± 0.0 | 0.1 ± 0.0 | 0.2 ± 0.0 | 0.1 ± 0.0 | 0.1 ± 0.0 | 0.2 ± 0.0 |
| Hesperitin | 1.2 ± 0.3 | 0.0 ± 0.0 | 0.0 ± 0.0 | 0.0 ± 0.0 | 0.0 ± 0.0 | 0.0 ± 0.0 | 0.0 ± 0.0 |
| Kaempferol | 72.9 ± 15.0 | 0.4 ± 0.0 | 0.8 ± 0.1 | 1.3 ± 0.1 | 0.6 ± 0.0 | 0.9 ± 0.0 | 1.6 ± 0.0 |
| Myricetin | 80.1 ± 20.0 | 25.1 ± 0.8 | 46.0 ± 0.4 | 71.1 ± 0.9 | 38.6 ± 0.5 | 48.4 ± 0.6 | 87.1 ± 0.8 |
| Quercetin-3-Glucoside | 5.2 ± 1.8 | 22.6 ± 0.7 | 28.2 ± 0.3 | 50.8 ± 0.7 | 27.1 ± 0.9 | 28.1 ± 0.5 | 55.3 ± 1.0 |
| Taxifolin | 1.4 ± 0.6 | 0.8 ± 0.1 | 1.0 ± 0.0 | 1.9 ± 0.1 | 1.0 ± 0.1 | 1.0 ± 0.0 | 2.0 ± 0.1 |
| Genstein | 0.0 ± 0.0 | 0.0 ± 0.0 | 0.1 ± 0.0 | 0.1 ± 0.0 | 0.1 ± 0.0 | 0.1 ± 0.0 | 0.1 ± 0.0 |
| Scopoletin | 0.1 ± 0.0 | 0.0 ± 0.0 | 0.1 ± 0.0 | 0.1 ± 0.0 | 0.1 ± 0.0 | 0.1 ± 0.0 | 0.1 ± 0.0 |
| Umbelliferone | 0.1 ± 0.1 | 0.0 ± 0.0 | 0.0 ± 0.0 | 0.0 ± 0.0 | 0.0 ± 0.0 | 0.0 ± 0.0 | 0.0 ± 0.0 |
| 7,8-dihydroxy-6-methyl coumarin | 36.4 ± 15.1 | 0.0 ± 0.0 | 0.1 ± 0.0 | 0.1 ± 0.0 | 0.1 ± 0.0 | 0.1 ± 0.0 | 0.1 ± 0.0 |
| Quercitrin | 0.3 ± 0.2 | 11.7 ± 0.6 | 25.4 ± 0.8 | 37.0 ± 1.0 | 17.0 ± 0.5 | 25.3 ± 0.3 | 42.3 ± 0.6 |
| Biochanin A | 0.0 ± 0.0 | 0.0 ± 0.0 | 0.1 ± 0.0 | 0.1 ± 0.0 | 0.1 ± 0.0 | 0.1 ± 0.0 | 0.1 ± 0.0 |
| Didymin | 8.8 ± 3.9 | 0.1 ± 0.0 | 0.3 ± 0.0 | 0.4 ± 0.0 | 0.3 ± 0.0 | 0.3 ± 0.0 | 0.6 ± 0.0 |
| Phloridzin | 0.0 ± 0.0 | 4.2 ± 0.2 | 5.6 ± 0.2 | 9.8 ± 0.3 | 4.8 ± 0.1 | 6.0 ± 0.1 | 10.8 ± 0.1 |
| Daidzein | 0.5 ± 0.1 | 0.0 ± 0.0 | 0.0 ± 0.0 | 0.0 ± 0.0 | 0.0 ± 0.0 | 0.0 ± 0.0 | 0.0 ± 0.0 |
| Luteolin | 4.5 ± 0.7 | 0.1 ± 0.0 | 0.3 ± 0.0 | 0.4 ± 0.0 | 0.2 ± 0.0 | 0.3 ± 0.0 | 0.5 ± 0.0 |
| Isorhamnetin | 0.0 ± 0.0 | 1.7 ± 0.1 | 3.7 ± 0.1 | 5.4 ± 0.1 | 2.6 ± 0.1 | 4.1 ± 0.1 | 6.7 ± 0.1 |
| Apigenin | 0.1 ± 0.0 | 0.1 ± 0.0 | 0.1 ± 0.0 | 0.2 ± 0.0 | 0.1 ± 0.0 | 0.1 ± 0.0 | 0.1 ± 0.0 |
| Gossypin | 19.1 ± 4.5 | 0.5 ± 0.0 | 0.9 ± 0.0 | 1.3 ± 0.0 | 0.7 ± 0.0 | 0.9 ± 0.0 | 1.6 ± 0.0 |
| Rutin | 0.0 ± 0.0 | 0.0 ± 0.0 | 0.0 ± 0.0 | 0.1 ± 0.0 | 0.1 ± 0.0 | 0.2 ± 0.0 | 0.3 ± 0.0 |
| **Benzoic Acids** | |  |  |  |  |  |  |
| salicylic acid | 15.0 ± 3.0 | 3.3 ± 0.1 | 4.5 ± 0.1 | 7.8 ± 0.1 | 2.2 ± 0.0 | 3.2 ± 0.1 | 5.4 ± 0.1 |
| m-hydroxybenzoic acid | 2.8 ± 1.1 | 0.1 ± 0.0 | 0.1 ± 0.0 | 0.2 ± 0.0 | 0.1 ± 0.0 | 0.1 ± 0.0 | 0.2 ± 0.0 |
| p-hydroxybenzoic acid | 20.6 ± 5.4 | 2.9 ± 0.2 | 3.5 ± 0.2 | 6.4 ± 0.3 | 2.6 ± 0.2 | 3.1 ± 0.1 | 5.7 ± 0.2 |
| 2,3-dihydroxybenzoic acid | 1.8 ± 0.8 | 0.0 ± 0.0 | 0.1 ± 0.0 | 0.1 ± 0.0 | 0.0 ± 0.0 | 0.1 ± 0.0 | 0.1 ± 0.0 |
| 2,4-dihydroxybenzoic acid | 0.0 ± 0.0 | 0.0 ± 0.0 | 0.0 ± 0.1 | 0.0 ± 0.1 | 0.0 ± 0.0 | 0.0 ± 0.0 | 0.0 ± 0.0 |
| 2,5-dihydroxybenzoic acid | 15.2 ± 6.7 | 0.0 ± 0.0 | 0.0 ± 0.0 | 0.0 ± 0.0 | 0.0 ± 0.0 | 0.0 ± 0.1 | 0.0 ± 0.1 |
| 2,6-dihydroxybenzoic acid | 0.0 ± 0.0 | 1.4 ± 0.0 | 1.8 ± 0.2 | 3.2 ± 0.2 | 1.8 ± 0.2 | 2.0 ± 0.0 | 3.9 ± 0.2 |
| protocatechuic acid | 188.2 ± 32.6 | 48.1 ± 1.3 | 59.7 ± 1.4 | 107.8 ± 1.9 | 54.5 ± 2.4 | 61.1 ± 2.7 | 115.5 ± 3.6 |
| p-anisic acid | 16.3 ± 3.1 | 5.0 ± 0.2 | 6.8 ± 0.5 | 11.8 ± 0.5 | 5.7 ± 0.3 | 8.7 ± 0.4 | 14.5 ± 0.5 |
| gallic acid | 74.0 ± 17.3 | 26.2 ± 0.5 | 34.1 ± 1.0 | 60.3 ± 1.2 | 35.0 ± 0.4 | 39.8 ± 1.5 | 74.8 ± 1.5 |
| vanillic acid | 72.6 ± 15.2 | 13.9 ± 0.3 | 15.1 ± 0.4 | 29.0 ± 0.5 | 17.3 ± 0.4 | 17.9 ± 0.6 | 35.1 ± 0.8 |
| syringic acid | 89.2 ± 31.8 | 34.0 ± 0.4 | 36.7 ± 0.9 | 70.7 ± 1.0 | 44.3 ± 1.2 | 45.4 ± 3.0 | 89.7 ± 3.2 |
| **Benzaldehydes** | |  |  |  |  |  |  |
| p-hydroxybenzaldehyde | 1.0 ± 0.1 | 0.2 ± 0.0 | 0.3 ± 0.0 | 0.5 ± 0.0 | 0.3 ± 0.0 | 0.3 ± 0.0 | 0.5 ± 0.0 |
| protocatachaldehyde | 14.3 ± 3.4 | 3.9 ± 0.3 | 4.0 ± 0.0 | 7.9 ± 0.3 | 3.4 ± 0.3 | 3.4 ± 0.0 | 6.8 ± 0.3 |
| 3,4,5-trihydroxybenzaldehyde | 11.4 ± 3.7 | 6.5 ± 0.1 | 6.8 ± 0.1 | 13.4 ± 0.1 | 6.5 ± 0.6 | 6.8 ± 0.1 | 13.3 ± 0.6 |
| vanillin | 0.6 ± 0.0 | 0.3 ± 0.0 | 0.3 ± 0.0 | 0.6 ± 0.0 | 0.3 ± 0.0 | 0.4 ± 0.0 | 0.7 ± 0.0 |
| syringin | 1.2 ± 0.2 | 0.8 ± 0.0 | 0.8 ± 0.0 | 1.6 ± 0.0 | 0.9 ± 0.0 | 0.9 ± 0.0 | 1.8 ± 0.0 |
| 3-methoxybenzaldehyde | 0.0 ± 0.0 | 0.0 ± 0.0 | 0.0 ± 0.0 | 0.0 ± 0.0 | 0.0 ± 0.0 | 0.0 ± 0.0 | 0.0 ± 0.0 |
| **Cinnamic Acids** | |  |  |  |  |  |  |
| cinnamic acid | 2.2 ± 0.8 | 1.6 ± 0.1 | 1.5 ± 0.1 | 3.1 ± 0.1 | 1.2 ± 0.1 | 1.1 ± 0.0 | 2.3 ± 0.1 |
| o-coumaric acid | 6.2 ± 2.7 | 0.0 ± 0.0 | 0.0 ± 0.0 | 0.0 ± 0.0 | 0.0 ± 0.0 | 0.0 ± 0.0 | 0.0 ± 0.0 |
| p-coumaric acid | 120.0 ± 38.8 | 63.3 ± 1.8 | 68.7 ± 2.3 | 132.0 ± 2.9 | 63.9 ± 4.0 | 67.2 ± 1.7 | 131.1 ± 4.3 |
| caffeic acid | 119.4 ± 28.4 | 20.6 ± 1.1 | 24.9 ± 0.4 | 45.5 ± 1.2 | 20.2 ± 1.2 | 22.7 ± 0.8 | 43.0 ± 1.5 |
| ferulic acid | 32.8 ± 5.6 | 6.4 ± 0.4 | 6.9 ± 0.2 | 13.3 ± 0.4 | 5.9 ± 0.5 | 6.2 ± 0.1 | 12.1 ± 0.5 |
| sinapic acid | 3.4 ± 0.7 | 2.2 ± 0.1 | 2.4 ± 0.2 | 4.6 ± 0.2 | 2.3 ± 0.1 | 2.5 ± 0.9 | 4.8 ± 0.9 |
| 3-methoxycinnamic acid | 0.0 ± 0.0 | 4.9 ± 0.3 | 4.8 ± 0.3 | 9.7 ± 0.4 | 4.4 ± 0.5 | 4.2 ± 0.1 | 8.6 ± 0.6 |
| 4-methoxycinnamic acid | 21.6 ± 9.0 | 0.3 ± 0.0 | 0.3 ± 0.0 | 0.6 ± 0.0 | 0.3 ± 0.0 | 0.3 ± 0.0 | 0.6 ± 0.0 |
| 3,4-dimethoxycinnamic acid | 0.8 ± 0.3 | 0.0 ± 0.0 | 0.0 ± 0.0 | 0.0 ± 0.0 | 0.0 ± 0.0 | 0.0 ± 0.0 | 0.0 ± 0.0 |
| 3,4,5-trimethoxycinnamic acid | 0.1 ± 0.0 | 0.1 ± 0.0 | 0.1 ± 0.0 | 0.1 ± 0.0 | 0.0 ± 0.0 | 0.0 ± 0.0 | 0.1 ± 0.0 |
| **Phenylpropionic acids** | |  |  |  |  |  |  |
| 2-hydroxyphenylpropionic acid | 0.1 ± 0.1 | 0.0 ± 0.0 | 0.0 ± 0.0 | 0.0 ± 0.0 | 0.0 ± 0.0 | 0.0 ± 0.0 | 0.0 ± 0.0 |
| 3-hydroxyphenylpropionic acid | 0.6 ± 0.3 | 0.0 ± 0.0 | 0.0 ± 0.0 | 0.0 ± 0.0 | 0.0 ± 0.0 | 0.0 ± 0.0 | 0.0 ± 0.0 |
| 4-hydroxyphenylpropionic acid | 2.6 ± 0.6 | 0.0 ± 0.0 | 0.0 ± 0.0 | 0.0 ± 0.0 | 0.0 ± 0.0 | 0.0 ± 0.0 | 0.0 ± 0.0 |
| 3,4-dihydroxyphenylpropionic acid | 2.1 ± 0.7 | 0.0 ± 0.0 | 0.0 ± 0.0 | 0.0 ± 0.0 | 0.0 ± 0.0 | 0.0 ± 0.0 | 0.0 ± 0.0 |
| 4-hydroxy-3-methoxyphenylpropionic acid | 2.7 ± 1.3 | 0.1 ± 0.0 | 0.1 ± 0.0 | 0.2 ± 0.0 | 0.1 ± 0.0 | 0.1 ± 0.0 | 0.1 ± 0.0 |
| **Benzenes** |  |  |  |  |  |  |  |
| 1,2-hydroxybenzene | 10.5 ± 3.9 | 0.4 ± 0.0 | 1.0 ± 0.0 | 1.3 ± 0.0 | 0.4 ± 0.0 | 0.9 ± 0.0 | 1.3 ± 0.0 |
| 1,2,3-trihydroxybenzene | 0.6 ± 0.3 | 0.1 ± 0.0 | 0.3 ± 0.0 | 0.4 ± 0.0 | 0.1 ± 0.0 | 0.3 ± 0.0 | 0.4 ± 0.0 |
| **Acetophenones** | |  |  |  |  |  |  |
| 4-hydroxyacetophenone | 0.4 ± 0.1 | 0.0 ± 0.0 | 0.0 ± 0.0 | 0.0 ± 0.0 | 0.0 ± 0.0 | 0.0 ± 0.0 | 0.0 ± 0.0 |
| 4-hydroxy-3-methoxyacetophenone | 0.3 ± 0.1 | 0.0 ± 0.0 | 0.0 ± 0.0 | 0.0 ± 0.0 | 0.0 ± 0.0 | 0.0 ± 0.0 | 0.0 ± 0.0 |
| 4-hydroxy-3,5-dimethoxyacetophenone | 0.0 ± 0.0 | 0.0 ± 0.0 | 0.0 ± 0.0 | 0.0 ± 0.0 | 0.0 ± 0.0 | 0.0 ± 0.0 | 0.1 ± 0.0 |
| **Phenylacetic acids** | |  |  |  |  |  |  |
| phenylacetic acid | 0.5 ± 0.1 | 0.4 ± 0.0 | 0.6 ± 0.0 | 0.9 ± 0.0 | 0.3 ± 0.0 | 0.6 ± 0.0 | 0.9 ± 0.0 |
| 4-hydroxyphenylacetic acid | 9.1 ± 3.6 | 0.1 ± 0.0 | 0.3 ± 0.0 | 0.4 ± 0.0 | 0.2 ± 0.0 | 0.3 ± 0.0 | 0.5 ± 0.0 |
| 3,4-dihydroxyphenylacetic acid | 2.9 ± 1.1 | 1.2 ± 0.1 | 1.2 ± 0.0 | 2.5 ± 0.1 | 0.4 ± 0.0 | 0.4 ± 0.1 | 0.7 ± 0.1 |
| **Mandelic Acids** | |  |  |  |  |  |  |
| 3-hydroxymandelic acid | 10.2 ± 4.4 | 2.2 ± 0.0 | 3.6 ± 0.1 | 5.8 ± 0.1 | 1.8 ± 0.1 | 2.9 ± 0.1 | 4.7 ± 0.1 |
| 4-hydroxymandelic acid | 0.0 ± 0.0 | 10.6 ± 0.8 | 12.1 ± 0.2 | 22.8 ± 0.8 | 13.3 ± 1.6 | 15.2 ± 1.2 | 28.5 ± 2.0 |
| 3,4-dihydroxymandelic acid | 2.1 ± 0.8 | 0.0 ± 0.0 | 0.0 ± 0.0 | 0.0. ± 0.0 | 0.0 ± 0.0 | 0.0 ± 0.0 | 0.0 ± 0.0 |
| **Phenyllactic Acids** | |  |  |  |  |  |  |
| 4-hydroxyphenyllactic acid | 13.0 ± 3.9 | 0.8 ± 0.0 | 2.2 ± 0.0 | 3.0 ± 0.1 | 0.8 ± 0.0 | 2.0 ± 0.1 | 2.8 ± 0.1 |
| **Phenolics Others** | |  |  |  |  |  |  |
| chlorogenic acid | 250.5 ± 118.0 | 52.1 ± 4.0 | 85.1 ± 1.0 | 137.1 ± 4.1 | 60.4 ± 0.3 | 71.9 ± 3.8 | 132.3 ± 3.8 |
| ethylferulate | 0.1 ± 0.1 | 0.1 ± 0.0 | 0.1 ± 0.0 | 0.2 ± 0.0 | 0.1 ± 0.0 | 0.1 ± 0.0 | 0.2 ± 0.0 |
| tyrosol | 5.0 ± 1.8 | 0.7 ± 0.0 | 1.7 ± 0.0 | 2.4 ± 0.1 | 0.6 ± 0.0 | 1.5 ± 0.1 | 2.1 ± 0.1 |
| Hydroxytyrosol | 25.0 ± 11.7 | 2.1 ± 0.1 | 2.4 ± 0.0 | 4.4 ± 0.1 | 1.5 ± 0.2 | 1.7 ± 0.0 | 3.2 ± 0.2 |
| **Phenolic dimers** | |  |  |  |  |  |  |
| Resveratrol | 0.1 ± 0.0 | 0.0 ± 0.0 | 0.0 ± 0.0 | 0.0 ± 0.0 | 0.0 ± 0.0 | 0.0 ± 0.0 | 0.0 ± 0.0 |
| **Indoles** |  |  |  |  |  |  |  |
| indole-3-acetic acid | 0.1 ± 0.0 | 0.1 ± 0.0 | 0.1 ± 0.0 | 0.1 ± 0.0 | 0.1 ± 0.0 | 0.1 ± 0.0 | 0.1 ± 0.0 |
| indole-3-carboxylic acid | 71.8 ± 21.9 | 0.1 ± 0.0 | 0.1 ± 0.0 | 0.2 ± 0.0 | 0.1 ± 0.0 | 0.1 ± 0.0 | 0.1 ± 0.0 |

Data are presented as mean ± standard deviation or sum ± root mean square.

Supplementary Table 4: Flavonoid and metabolite content of grape seed extract

| GRAPE SEED |  |  |  |  |  |  |  |
| --- | --- | --- | --- | --- | --- | --- | --- |
| ng/100mg extract | **raw extract** | **gastric phase** | **gastric phase** | **gastric phase** | **intestinal phase** | **intestinal phase** | **intestinal phase** |
| **Flavanoids/Coumarins** | | **pellet** | **supernatant** | **sum** | **pellet** | **supernatant** | **sum** |
| Catechin | 1074.6 ± 325.1 | 264.8 ± 22.4 | 344.8 ± 4.4 | 609.6 ± 22.8 | 267.5 ± 15.0 | 341.0 ± 6.2 | 608.5 ± 16.3 |
| Epicatechin | 554.2 ± 209.6 | 121.2 ± 3.4 | 125.8 ± 3.0 | 247.0 ± 4.5 | 117.6 ± 2.4 | 123.2 ± 5.5 | 240.7 ± 6.0 |
| Gallocatechin | 43.4 ± 19.1 | 12.9 ± 0.3 | 7.2 ± 0.5 | 20.0 ± 0.6 | 13.1 ± 0.1 | 7.2 ± 0.3 | 20.3 ± 0.4 |
| Epigallocatechin | 1.4 ± 0.7 | 21.8 ± 0.0 | 12.8 ± 0.0 | 34.7 ± 0.0 | 20.5 ± 0.0 | 14.4 ± 0.0 | 34.8 ± 0.0 |
| Epigallocatechin Gallate | 20.7 ± 5.5 | 1.1 ± 0.3 | 2.9 ± 0.3 | 4.0 ± 0.4 | 1.4 ± 0.3 | 3.1 ± 0.1 | 4.5 ± 0.3 |
| Quercetin | 21.5 ± 7.0 | 19.4 ± 1.6 | 17.7 ± 0.8 | 37.1 ± 1.8 | 23.9 ± 0.3 | 21.8 ± 0.5 | 45.7 ± 0.6 |
| Isoliquiritigenin | 0.0 ± 0.0 | 0.0 ± 0.0 | 0.0 ± 0.0 | 0.0 ± 0.0 | 0.0 ± 0.0 | 0.0 ± 0.0 | 0.0 ± 0.0 |
| Phloretin | 0.3 ± 0.1 | 0.1 ± 0.0 | 0.2 ± 0.0 | 0.3 ± 0.0 | 0.2 ± 0.0 | 0.2 ± 0.0 | 0.4 ± 0.0 |
| Naringenin | 1.5 ± 0.7 | 2.7 ± 0.2 | 2.8 ± 0.3 | 5.6 ± 0.4 | 2.8 ± 0.0 | 3.0 ± 0.0 | 5.8 ± 0.1 |
| Naringin | 0.0 ± 0.0 | 0.1 ± 0.0 | 0.1 ± 0.0 | 0.2 ± 0.0 | 0.1 ± 0.0 | 0.1 ± 0.0 | 0.2 ± 0.0 |
| Hesperitin | 0.0 ± 0.0 | 0.0 ± 0.0 | 0.0 ± 0.0 | 0.0 ± 0.0 | 0.0 ± 0.0 | 0.0 ± 0.0 | 0.1 ± 0.0 |
| Kaempferol | 2.3 ± 1.0 | 1.2 ± 0.2 | 1.2 ± 0.3 | 2.4 ± 0.3 | 1.8 ± 0.1 | 1.9 ± 0.0 | 3.7 ± 0.1 |
| Myricetin | 0.9 ± 0.4 | 27.1 ± 0.1 | 22.9 ± 0.1 | 50.1 ± 0.1 | 35.3 ± 0.0 | 31.3 ± 0.0 | 66.6 ± 0.0 |
| Quercetin-3-Glucoside | 6.0 ± 2.6 | 29.4 ± 0.1 | 22.6 ± 0.1 | 52.1 ± 0.1 | 29.5 ± 0.1 | 24.4 ± 0.1 | 53.9 ± 0.2 |
| Taxifolin | 28.6 ± 10.6 | 2.9 ± 0.6 | 4.9 ± 0.4 | 7.8 ± 0.7 | 3.2 ± 0.3 | 4.7 ± 0.1 | 7.9 ± 0.3 |
| Genstein | 0.0 ± 0.0 | 0.1 ± 0.0 | 0.1 ± 0.0 | 0.1 ± 0.0 | 0.1 ± 0.0 | 0.1 ± 0.0 | 0.2 ± 0.0 |
| Scopoletin | 0.0 ± 0.0 | 0.0 ± 0.0 | 0.0 ± 0.0 | 0.1 ± 0.0 | 0.0 ± 0.0 | 0.0 ± 0.0 | 0.1 ± 0.0 |
| 7,8-dihydroxy-6-methyl coumarin | 0.0 ± 0.0 | 0.1 ± 0.0 | 0.1 ± 0.0 | 0.1 ± 0.0 | 0.1 ± 0.0 | 0.1 ± 0.0 | 0.2 ± 0.0 |
| Quercitrin | 0.7 ± 0.3 | 7.4 ± 0.1 | 7.0 ± 0.0 | 14.3 ± 0.1 | 8.9 ± 0.0 | 8.8 ± 0.0 | 17.7 ± 0.0 |
| Biochanin A | 0.0 ± 0.0 | 0.0 ± 0.0 | 0.0 ± 0.0 | 0.1 ± 0.0 | 0.1 ± 0.0 | 0.1 ± 0.0 | 0.1 ± 0.0 |
| Didymin | 0.0 ± 0.0 | 0.1 ± 0.0 | 0.1 ± 0.0 | 0.2 ± 0.0 | 0.2 ± 0.0 | 0.2 ± 0.0 | 0.3 ± 0.0 |
| Phloridzin | 1.8 ± 0.8 | 1.9 ± 0.0 | 2.0 ± 0.0 | 3.9 ± 0.1 | 1.8 ± 0.0 | 1.9 ± 0.0 | 3.7 ± 0.0 |
| Luteolin | 0.5 ± 0.2 | 0.2 ± 0.1 | 0.2 ± 0.0 | 0.4 ± 0.1 | 0.3 ± 0.0 | 0.4 ± 0.0 | 0.7 ± 0.0 |
| Isorhamnetin | 0.9 ± 0.4 | 3.5 ± 0.2 | 1.8 ± 0.1 | 5.3 ± 0.2 | 3.8 ± 0.0 | 2.6 ± 0.0 | 6.5 ± 0.0 |
| Apigenin | 0.1 ± 0.0 | 0.1 ± 0.0 | 0.1 ± 0.0 | 0.1 ± 0.0 | 0.1 ± 0.0 | 0.1 ± 0.0 | 0.2 ± 0.0 |
| Gossypin | 0.5 ± 0.2 | 0.5 ± 0.0 | 0.3 ± 0.0 | 0.9 ± 0.0 | 0.6 ± 0.0 | 0.4 ± 0.0 | 1.0 ± 0.0 |
| Rutin | 0.0 ± 0.0 | 0.3 ± 0.0 | 0.2 ± 0.0 | 0.5 ± 0.0 | 0.4 ± 0.0 | 0.2 ± 0.0 | 0.6 ± 0.0 |
| **Benzoic Acids** | |  |  |  |  |  |  |
| salicylic acid | 0.2 ± 0.1 | 2.6 ± 0.0 | 2.0 ± 0.0 | 4.6 ± 0.0 | 2.0 ± 0.0 | 1.6 ± 0.0 | 3.6 ± 0.0 |
| m-hydroxybenzoic acid | 0.0 ± 0.0 | 0.2 ± 0.0 | 0.1 ± 0.0 | 0.2 ± 0.0 | 0.2 ± 0.0 | 0.1 ± 0.0 | 0.2 ± 0.0 |
| p-hydroxybenzoic acid | 3.1 ± 0.6 | 4.9 ± 0.0 | 2.1 ± 0.0 | 6.9 ± 0.1 | 3.9 ± 0.1 | 2.4 ± 0.0 | 6.2 ± 0.1 |
| 2,3-dihydroxybenzoic acid | 0.0 ± 0.0 | 0.1 ± 0.0 | 0.1 ± 0.0 | 0.2 ± 0.0 | 0.1 ± 0.0 | 0.1 ± 0.0 | 0.1 ± 0.0 |
| 2,4-dihydroxybenzoic acid | 0.1 ± 0.0 | 0.0 ± 0.0 | 0.0 ± 0.0 | 0.0 ± 0.0 | 0.0 ± 0.0 | 0.0 ± 0.0 | 0.0 ± 0.0 |
| 2,5-dihydroxybenzoic acid | 0.0 ± 0.0 | 0.7 ± 0.0 | 0.7 ± 0.0 | 1.4 ± 0.0 | 0.4 ± 0.0 | 0.4 ± 0.0 | 0.8 ± 0.0 |
| 2,6-dihydroxybenzoic acid | 0.0 ± 0.0 | 3.5 ± 0.0 | 0.8 ± 0.0 | 4.3 ± 0.0 | 2.6 ± 0.0 | 1.0 ± 0.0 | 3.6 ± 0.0 |
| protocatechuic acid | 45.0 ± 14.6 | 59.2 ± 1.3 | 41.7 ± 0.5 | 100.9 ± 1.4 | 55 ± 0.2 | 48.5 ± 1.6 | 103.5 ± 1.6 |
| p-anisic acid | 0.9 ± 0.2 | 8.0 ± 0.0 | 4.7 ± 0.0 | 12.7 ± 0.0 | 7.8 ± 0.1 | 5.1 ± 0.0 | 12.9 ± 0.1 |
| gallic acid | 166.8 ± 52.0 | 42.2 ± 1.6 | 45.1 ± 2.3 | 87.3 ± 2.8 | 42.8 ± 0.5 | 53.9 ± 2.2 | 96.7 ± 2.3 |
| vanillic acid | 1.9 ± 0.3 | 25.1 ± 0.0 | 10.2 ± 0.1 | 35.2 ± 0.1 | 20.8 ± 0.1 | 13.4 ± 0.0 | 34.3 ± 0.1 |
| syringic acid | 1.3 ± 0.3 | 69.4 ± 0.1 | 26.3 ± 0.1 | 95.7 ± 0.1 | 57.3 ± 0.1 | 36.6 ± 0.1 | 93.8 ± 0.1 |
| **Benzaldehydes** | |  |  |  |  |  |  |
| p-hydroxybenzaldehyde | 1.1 ± 0.3 | 0.2 ± 0.0 | 0.3 ± 0.0 | 0.5 ± 0.0 | 0.1 ± 0.0 | 0.2 ± 0.0 | 0.4 ± 0.0 |
| protocatachaldehyde | 85.2 ± 23.2 | 1.5 ± 0.5 | 5.6 ± 0.3 | 7.1 ± 0.5 | 1.1 ± 0.5 | 6.3 ± 0.5 | 7.5 ± 0.7 |
| 3,4,5-trihydroxybenzaldehyde | 2.6 ± 1.2 | 3.3 ± 0.1 | 2.6 ± 0.1 | 5.9 ± 0.1 | 1.3 ± 0.1 | 3.2 ± 0.0 | 4.5 ± 0.1 |
| vanillin | 4.3 ± 1.4 | 1.6 ± 0.3 | 2.1 ± 0.1 | 3.7 ± 0.3 | 1.4 ± 0.1 | 2.0 ± 0.0 | 3.4 ± 0.1 |
| syringin | 0.1 ± 0.0 | 0.4 ± 0.0 | 0.3 ± 0.0 | 0.7 ± 0.0 | 0.3 ± 0.0 | 0.4 ± 0.0 | 0.7 ± 0.0 |
| 3-methoxybenzaldehyde | 0.1 ± 0.0 | 0.0 ± 0.0 | 0.0 ± 0.0 | 0.0 ± 0.0 | 0.0 ± 0.0 | 0.0 ± 0.0 | 0.0 ± 0.0 |
| **Cinnamic Acids** | |  |  |  |  |  |  |
| cinnamic acid | 1.8 ± 0.5 | 2.4 ± 0.1 | 2.0 ± 0.1 | 4.4 ± 0.1 | 2.0 ± 0.1 | 1.7 ± 0.1 | 3.7 ± 0.1 |
| o-coumaric acid | 0.0 ± 0.0 | 0.0 ± 0.0 | 0.0 ± 0.0 | 0.0 ± 0.0 | 0.0 ± 0.0 | 0.0 ± 0.0 | 0.0 ± 0.0 |
| p-coumaric acid | 4.3 ± 1.1 | 88.6 ± 0.1 | 36.4 ± 0.1 | 125.0 ± 0.2 | 74.0 ± 0.0 | 42.9 ± 0.1 | 116.9 ± 0.1 |
| caffeic acid | 0.9 ± 0.2 | 22.2 ± 0.0 | 14.8 ± 0.0 | 37.1 ± 0.0 | 19.9 ± 0.0 | 14.8 ± 0.0 | 34.7 ± 0.0 |
| ferulic acid | 15.4 ± 3.7 | 8.0 ± 0.2 | 8.2 ± 0.2 | 16.2 ± 0.3 | 6.8 ± 0.1 | 8.7 ± 0.2 | 15.5 ± 0.2 |
| sinapic acid | 0.2 ± 0.0 | 7.4 ± 0.0 | 6.7 ± 0.0 | 14.1 ± 0.0 | 7.3 ± 0.0 | 6.7 ± 0.0 | 14.1 ± 0.0 |
| 3-methoxycinnamic acid | 0.0 ± 0.0 | 7.5 ± 0.0 | 3.4 ± 0.0 | 10.9 ± 0.0 | 5.6 ± 0.0 | 3.5 ± 0.0 | 9.1 ± 0.0 |
| 4-methoxycinnamic acid | 0.2 ± 0.1 | 0.6 ± 0.0 | 0.2 ± 0.0 | 0.8 ± 0.0 | 0.4 ± 0.0 | 0.2 ± 0.0 | 0.7 ± 0.0 |
| 3,4,5-trimethoxycinnamic acid | 0.1 ± 0.0 | 0.1 ± 0.0 | 0.1 ± 0.0 | 0.2 ± 0.0 | 0.1 ± 0.0 | 0.1 ± 0.0 | 0.2 ± 0.0 |
| **Phenylpropionic acids** | |  |  |  |  |  |  |
| phenylpropionic acid | 0.5 ± 0.3 | 0.1 ± 0.0 | 0.1 ± 0.1 | 0.3 ± 0.1 | 0.1 ± 0.0 | 0.1 ± 0.0 | 0.2 ± 0.0 |
| 4-hydroxyphenylpropionic acid | 0.0 ± 0.0 | 0.0 ± 0.0 | 0.0 ± 0.0 | 0.0 ± 0.0 | 0.0 ± 0.0 | 0.0 ± 0.1 | 0.0 ± 0.1 |
| 4-hydroxy-3-methoxyphenylpropionic acid | 0.8 ± 0.4 | 0.3 ± 0.0 | 0.3 ± 0.0 | 0.7 ± 0.0 | 0.3 ± 0.0 | 0.3 ± 0.0 | 0.6 ± 0.0 |
| **Benzenes** |  |  |  |  |  |  |  |
| 1,2-hydroxybenzene | 5.0 ± 2.4 | 0.2 ± 0.0 | 0.2 ± 0.0 | 0.3 ± 0.0 | 0.1 ± 0.0 | 0.1 ± 0.0 | 0.3 ± 0.0 |
| 1,2,3-trihydroxybenzene | 0.0 ± 0.0 | 0.1 ± 0.0 | 0.1 ± 0.0 | 0.2 ± 0.0 | 0.1 ± 0.0 | 0.1 ± 0.0 | 0.2 ± 0.0 |
| **Acetophenones** | |  |  |  |  |  |  |
| 4-hydroxyacetophenone | 0.1 ± 0.0 | 0.0 ± 0.0 | 0.0 ± 0.0 | 0.0 ± 0.0 | 0.0 ± 0.0 | 0.0 ± 0.0 | 0.0 ± 0.0 |
| 4-hydroxy-3-methoxyacetophenone | 0.3 ± 0.1 | 0.2 ± 0.0 | 0.2 ± 0.0 | 0.3 ± 0.0 | 0.1 ± 0.0 | 0.2 ± 0.0 | 0.3 ± 0.0 |
| 4-hydroxy-3,5-dimethoxyacetophenone | 0.0 ± 0.0 | 0.1 ± 0.0 | 0.0 ± 0.0 | 0.1 ± 0.0 | 0.1 ± 0.0 | 0.0 ± 0.0 | 0.1 ± 0.0 |
| **Phenylacetic acids** | |  |  |  |  |  |  |
| phenylacetic acid | 0.2 ± 0.1 | 0.3 ± 0.0 | 0.3 ± 0.0 | 0.6 ± 0.0 | 0.3 ± 0.0 | 0.2 ± 0.0 | 0.5 ± 0.0 |
| 4-hydroxyphenylacetic acid | 1.3 ± 0.6 | 0.1 ± 0.0 | 0.1 ± 0.0 | 0.2 ± 0.0 | 0.1 ± 0.0 | 0.1 ± 0.0 | 0.2 ± 0.0 |
| 3,4-dihydroxyphenylacetic acid | 41.1 ± 18.6 | 0.5 ± 1.2 | 2.3 ± 0.6 | 2.8 ± 1.4 | 2.3 ± 0.9 | 1.5 ± 1.1 | 3.8 ± 1.4 |
| **Mandelic Acids** | |  |  |  |  |  |  |
| 3-hydroxymandelic acid | 1.1 ± 0.3 | 1.7 ± 0.8 | 4.7 ± 0.2 | 6.4 ± 0.8 | 1.8 ± 0.1 | 4.7 ± 0.1 | 6.4 ± 0.1 |
| 4-hydroxymandelic acid | 3.3 ± 1.5 | 24.7 ± 0.6 | 28.5 ± 2.0 | 53.3 ± 2.1 | 20.8 ± 0.9 | 36.9 ± 0.7 | 57.7 ± 1.2 |
| 3,4-dihydroxymandelic acid | 6.4 ± 3.0 | 0.0 ± 0.0 | 0.0 ± 0.0 | 0.0 ± 0.0 | 0.0 ± 0.0 | 0.0 ± 0.0 | 0.0 ± 0.0 |
| **Phenyllactic Acids** | |  |  |  |  |  |  |
| 4-hydroxyphenyllactic acid | 0.5 ± 0.2 | 0.9 ± 0.0 | 0.8 ± 0.1 | 1.7 ± 0.1 | 0.9 ± 0.0 | 0.9 ± 0.0 | 1.8 ± 0.0 |
| **Phenolics Others** | |  |  |  |  |  |  |
| chlorogenic acid | 0.0 ± 0.0 | 38.0 ± 0.0 | 38.0 ± 0.0 | 76.0 ± 0.0 | 40.7 ± 0.0 | 40.7 ± 0.0 | 81.5 ± 0.0 |
| ethylferulate | 0.5 ± 0.2 | 0.2 ± 0.0 | 0.2 ± 0.0 | 0.5 ± 0.0 | 0.3 ± 0.0 | 0.3 ± 0.0 | 0.5 ± 0.0 |
| tyrosol | 0.6 ± 0.3 | 0.7 ± 0.0 | 0.7 ± 0.0 | 1.4 ± 0.0 | 0.6 ± 0.0 | 0.6 ± 0.1 | 1.3 ± 0.1 |
| Hydroxytyrosol | 0.0 ± 0.0 | 0.8 ± 0.0 | 1.5 ± 0.0 | 2.3 ± 0.0 | 0.5 ± 0.0 | 1.1 ± 0.0 | 1.6 ± 0.0 |
| **Phenolic dimers** | |  |  |  |  |  |  |
| Resveratrol | 3.1 ± 1.4 | 1.2 ± 0.3 | 1.4 ± 0.1 | 2.6 ± 0.3 | 1.5 ± 0.1 | 1.5 ± 0.0 | 3.0 ± 0.1 |
| **Indoles** |  |  |  |  |  |  |  |
| indole-3-acetic acid | 0.1 ± 0.1 | 0.1 ± 0.0 | 0.1 ± 0.0 | 0.2 ± 0.0 | 0.1 ± 0.0 | 0.1 ± 0.0 | 0.2 ± 0.0 |
| indole-3-carboxylic acid | 20.5 ± 5.6 | 0.1 ± 0.0 | 0.0 ± 0.0 | 0.1 ± 0.0 | 0.1 ± 0.0 | 0.0 ± 0.0 | 0.1 ± 0.0 |

Data are presented as mean ± standard deviation or sum ± root mean square, and rounded up.

Supplementary Table 5: Flavonoid and metabolite content in the bilberry-grape seed formulation extract

| FORMULATION | **raw extract** | **gastric phase** | **gastric phase** | **gastric phase** | **intestinal phase** | **intestinal phase** | **intestinal phase** |
| --- | --- | --- | --- | --- | --- | --- | --- |
| ng/100 mg extract |  | **Pellet** | **supernatant** | **sum** | **pellet** | **supernatant** | **sum** |
| **Flavanoids/Coumarins** |  |  |  |  |  |  |  |
| Catechin | 300.5 ± 12.1 | 60.9 ± 7.3 | 57.8 ± 11.8 | 118.8 ± 13.8 | 56.7 ± 22.5 | 52.7 ± 9.2 | 109.5 ± 24.3 |
| Epicatechin | 132.6 ± 9.0 | 82.1 ± 0.5 | 78.0 ± 2.6 | 160.1 ± 2.6 | 83.3 ± 7.0 | 78.1 ± 1.7 | 161.4 ± 7.2 |
| Gallocatechin | 10.0 ± 1.0 | 35.1 ± 0.7 | 35.0 ± 4.2 | 70.1 ± 4.3 | 32.2 ± 0.9 | 31.8 ± 0.8 | 64.0 ± 1.2 |
| Epigallocatechin | 17.6 ± 2.0 | 60.1 ± 1.2 | 59.7 ± 1.8 | 119.8 ± 2.1 | 54.3 ± 1.6 | 53.2 ± 0.5 | 107.5 ± 1.7 |
| Epigallocatechin Gallate | 2.3 ± 0.3 | 0.0 ± 0.1 | 0.0 ± 0.3 | 0.0 ± 0.4 | 0.0 ± 0.4 | 0.0 ± 0.1 | 0.0 ± 0.5 |
| Quercetin | 24.5 ± 3.1 | 24.9 ± 0.6 | 27.8 ± 1.2 | 52.6 ± 1.3 | 25.4 ± 2.2 | 27.6 ± 0.5 | 53.0 ± 2.3 |
| Isoliquiritigenin | 0.0 ± 0.0 | 0.0 ± 0.0 | 0.0 ± 0.0 | 0.1 ± 0.0 | 0.0 ± 0.0 | 0.0 ± 0.0 | 0.1 ± 0.0 |
| Phloretin | 0.3 ± 0.1 | 0.1 ± 0.0 | 0.1 ± 0.0 | 0.2 ± 0.0 | 0.1 ± 0.0 | 0.1 ± 0.0 | 0.2 ± 0.0 |
| Naringenin | 2.4 ± 0.4 | 2.9 ± 0.2 | 3.0 ± 0.3 | 5.8 ± 0.4 | 2.4 ± 0.5 | 2.4 ± 0.0 | 4.8 ± 0.5 |
| Naringin | 0.0 ± 0.0 | 0.2 ± 0.0 | 0.2 ± 0.0 | 0.3 ± 0.0 | 0.2 ± 0.0 | 0.2 ± 0.0 | 0.3 ± 0.0 |
| Hesperitin | 0.0 ± 0.0 | 0.0 ± 0.0 | 0.0 ± 0.0 | 0.0 ± 0.0 | 0.0 ± 0.0 | 0.0 ± 0.0 | 0.0 ± 0.0 |
| Kaempferol | 1.7 ± 0.4 | 0.9 ± 0.1 | 0.9 ± 0.2 | 1.8 ± 0.3 | 0.9 ± 0.2 | 1.0 ± 0.1 | 1.9 ± 0.2 |
| Myricetin | 51.0 ± 6.5 | 48.5 ± 0.8 | 52.6 ± 2.2 | 101.1 ± 2.3 | 51.3 ± 3.1 | 54.7 ± 0.8 | 106.0 ± 3.2 |
| Quercetin-3-Glucoside | 31.4 ± 1.9 | 31.4 ± 0.4 | 35.6 ± 0.8 | 67.0 ± 1.0 | 30.4 ± 1.0 | 34.0 ± 0.2 | 64.4 ± 1.1 |
| Taxifolin | 5.7 ± 0.9 | 0.6 ± 0.3 | 0.6 ± 0.3 | 1.3 ± 0.4 | 0.9 ± 0.5 | 0.9 ± 0.2 | 1.8 ± 0.5 |
| Genstein | 0.0 ± 0.0 | 0.1 ± 0.0 | 0.1 ± 0.0 | 0.2 ± 0.0 | 0.1 ± 0.0 | 0.1 ± 0.0 | 0.2 ± 0.0 |
| Scopoletin | 0.8 ± 0.0 | 0.1 ± 0.0 | 0.1 ± 0.0 | 0.1 ± 0.0 | 0.1 ± 0.0 | 0.1 ± 0.0 | 0.1 ± 0.0 |
| Umbelliferone | 0.0 ± 0.0 | 0.0 ± 0.0 | 0.0 ± 0.0 | 0.0 ± 0.0 | 0.0 ± 0.0 | 0.0 ± 0.0 | 0.0 ± 0.0 |
| 7,8-dihydroxy-6-methyl coumarin | 0.1 ± 0.0 | 0.1 ± 0.0 | 0.1 ± 0.0 | 0.2 ± 0.0 | 0.1 ± 0.0 | 0.1 ± 0.0 | 0.2 ± 0.0 |
| Quercitrin | 10.2 ± 1.3 | 24.7 ± 0.4 | 24.7 ± 1.2 | 49.4 ± 1.2 | 25.4 ± 0.5 | 25.3 ± 0.8 | 50.7 ± 0.9 |
| Biochanin A | 0.0 ± 0.0 | 0.1 ± 0.0 | 0.1 ± 0.0 | 0.2 ± 0.0 | 0.1 ± 0.0 | 0.1 ± 0.0 | 0.1 ± 0.0 |
| Didymin | 0.0 ± 0.0 | 0.3 ± 0.0 | 0.3 ± 0.0 | 0.6 ± 0.0 | 0.3 ± 0.0 | 0.3 ± 0.0 | 0.6 ± 0.0 |
| Phloridzin | 3.2 ± 0.4 | 5.0 ± 0.1 | 5.0 ± 0.2 | 10.0 ± 0.2 | 5.9 ± 0.1 | 5.9 ± 0.0 | 11.8 ± 0.1 |
| Luteolin | 0.8 ± 0.1 | 0.3 ± 0.0 | 0.3 ± 0.1 | 0.6 ± 0.1 | 0.3 ± 0.0 | 0.3 ± 0.0 | 0.5 ± 0.0 |
| Isorhamnetin | 1.9 ± 0.4 | 5.1 ± 0.0 | 5.3 ± 0.4 | 10.4 ± 0.4 | 5.1 ± 0.3 | 5.3 ± 0.0 | 10.4 ± 0.3 |
| Formononetin | 0.0 ± 0.0 | 0.0 ± 0.0 | 0.0 ± 0.0 | 0.0 ± 0.0 | 0.0 ± 0.0 | 0.0 ± 0.0 | 0.0 ± 0.0 |
| Apigenin | 0.1 ± 0.0 | 0.1 ± 0.0 | 0.1 ± 0.0 | 0.2 ± 0.0 | 0.1 ± 0.0 | 0.1 ± 0.0 | 0.2 ± 0.0 |
| Gossypin | 5.8 ± 0.6 | 1.0 ± 0.0 | 1.1 ± 0.0 | 2.1 ± 0.0 | 1.0 ± 0.0 | 1.1 ± 0.0 | 2.1 ± 0.0 |
| Rutin | 0.0 ± 0.0 | 0.1 ± 0.0 | 0.2 ± 0.0 | 0.3 ± 0.0 | 0.4 ± 0.0 | 0.4 ± 0.0 | 0.7 ± 0.0 |
| **Benzoic Acids** |  |  |  |  |  |  |  |
| salicylic acid | 5.0 ± 0.6 | 4.7 ± 0.1 | 5.2 ± 0.1 | 9.9 ± 0.1 | 3.5 ± 0.1 | 4.0 ± 0.1 | 7.5 ± 0.1 |
| m-hydroxybenzoic acid | 0.8 ± 0.1 | 0.2 ± 0.0 | 0.2 ± 0.0 | 0.4 ± 0.0 | 0.2 ± 0.0 | 0.2 ± 0.0 | 0.4 ± 0.0 |
| p-hydroxybenzoic acid | 7.5 ± 0.5 | 5.6 ± 0.1 | 5.7 ± 0.1 | 11.3 ± 0.1 | 4.6 ± 0.1 | 4.7 ± 0.0 | 9.3 ± 0.1 |
| 2,3-dihydroxybenzoic acid | 0.4 ± 0.0 | 0.1 ± 0.0 | 0.1 ± 0.0 | 0.2 ± 0.0 | 0.1 ± 0.0 | 0.1 ± 0.0 | 0.2 ± 0.0 |
| 2,5-dihydroxybenzoic acid | 4.3 ± 0.2 | 0.6 ± 0.0 | 1.1 ± 0.0 | 1.8 ± 0.0 | 0.6 ± 0.0 | 0.9 ± 0.0 | 1.5 ± 0.0 |
| 2,6-dihydroxybenzoic acid | 0.7 ± 0.1 | 4.0 ± 0.1 | 4.1 ± 0.0 | 8.1 ± 0.1 | 3.1 ± 0.1 | 3.2 ± 0.1 | 6.3 ± 0.1 |
| protocatechuic acid | 86.6 ± 7.6 | 72.3 ± 0.6 | 76.3 ± 1.8 | 148.6 ± 1.9 | 68.4 ± 0.8 | 70.3 ± 0.5 | 138.7 ± 0.9 |
| p-anisic acid | 2.6 ± 0.2 | 8.6 ± 0.2 | 10.1 ± 0.3 | 18.7 ± 0.3 | 11.1 ± 0.4 | 11.9 ± 0.2 | 23.0 ± 0.5 |
| gallic acid | 192.2 ± 8.9 | 47.2 ± 0.8 | 50.2 ± 2.0 | 97.4 ± 2.1 | 47.1 ± 1.0 | 49.2 ± 0.6 | 96.2 ± 1.2 |
| vanillic acid | 34.1 ± 1.9 | 25.7 ± 0.1 | 27.1 ± 0.3 | 52.8 ± 0.4 | 22.2 ± 0.4 | 23.0 ± 0.2 | 45.2 ± 0.4 |
| syringic acid | 62.0 ± 4.9 | 69.4 ± 0.2 | 73.8 ± 0.8 | 143.2 ± 0.8 | 59.2 ± 1.7 | 61.8 ± 0.8 | 121.0 ± 1.8 |
| **Benzaldehydes** |  |  |  |  |  |  |  |
| p-hydroxybenzaldehyde | 0.5 ± 0.1 | 0.2 ± 0.0 | 0.2 ± 0.0 | 0.3 ± 0.0 | 0.1 ± 0.0 | 0.1 ± 0.0 | 0.2 ± 0.0 |
| protocatachaldehyde | 29.7 ± 1.5 | 1.4 ± 0.2 | 1.3 ± 0.4 | 2.7 ± 0.4 | 1.0 ± 0.6 | 0.8 ± 0.0 | 1.8 ± 0.6 |
| 3,4,5-trihydroxybenzaldehyde | 4.2 ± 1.0 | 3.8 ± 0.3 | 3.6 ± 0.1 | 7.5 ± 0.4 | 2.0 ± 0.0 | 1.7 ± 0.1 | 3.6 ± 0.1 |
| vanillin | 2.0 ± 0.3 | 0.2 ± 0.1 | 0.2 ± 0.0 | 0.5 ± 0.1 | 0.2 ± 0.1 | 0.2 ± 0.0 | 0.4 ± 0.1 |
| syringin | 0.2 ± 0.0 | 0.6 ± 0.0 | 0.6 ± 0.0 | 1.2 ± 0.0 | 0.5 ± 0.0 | 0.5 ± 0.0 | 1.0 ± 0.0 |
| 20HBAldc | 0.1 ± 0.0 | 0.0 ± 0.0 | 0.0 ± 0.0 | 0.0 ± 0.0 | 0.0 ± 0.0 | 0.1 ± 0.0 | 0.1 ± 0.0 |
| **Cinnamic Acids** |  |  |  |  |  |  |  |
| cinnamic acid | 2.1 ± 0.3 | 1.9 ± 0.2 | 1.9 ± 0.1 | 3.8 ± 0.2 | 1.6 ± 0.2 | 1.6 ± 0.0 | 3.1 ± 0.2 |
| o-coumaric acid | 1.3 ± 0.0 | 0.0 ± 0.0 | 0.0 ± 0.0 | 0.0 ± 0.0 | 0.0 ± 0.0 | 0.0 ± 0.0 | 0.0 ± 0.0 |
| p-coumaric acid | 88.2 ± 4.4 | 99.7 ± 0.7 | 99.1 ± 1.6 | 198.8 ± 1.7 | 86.1 ± 0.6 | 84.6 ± 0.8 | 170.7 ± 0.9 |
| caffeic acid | 63.6 ± 4.6 | 28.8 ± 0.4 | 29.8 ± 0.5 | 58.6 ± 0.6 | 26.4 ± 0.5 | 27.1 ± 0.2 | 53.5 ± 0.5 |
| ferulic acid | 7.1 ± 0.5 | 6.9 ± 0.4 | 7.7 ± 0.3 | 14.7 ± 0.4 | 5.9 ± 0.4 | 6.5 ± 0.0 | 12.5 ± 0.4 |
| sinapic acid | 1.3 ± 0.1 | 2.8 ± 0.0 | 7.7 ± 0.0 | 10.5 ± 0.0 | 3.0 ± 0.0 | 7.7 ± 0.1 | 10.8 ± 0.1 |
| 3-methoxycinnamic acid | 0.0 ± 0.0 | 7.7 ± 0.3 | 7.6 ± 0.3 | 15.3 ± 0.5 | 5.8 ± 0.3 | 5.7 ± 0.1 | 11.5 ± 0.3 |
| 4-methoxycinnamic acid | 5.7 ± 0.4 | 0.6 ± 0.0 | 0.6 ± 0.0 | 1.3 ± 0.0 | 0.4 ± 0.0 | 0.4 ± 0.0 | 0.9 ± 0.0 |
| 3,4-dimethoxycinnamic acid | 0.3 ± 0.0 | 0.0 ± 0.0 | 0.0 ± 0.0 | 0.0 ± 0.0 | 0.0 ± 0.0 | 0.0 ± 0.0 | 0.0 ± 0.0 |
| 3,4,5-trimethoxycinnamic acid | 0.1 ± 0.0 | 0.1 ± 0.0 | 0.1 ± 0.0 | 0.2 ± 0.0 | 0.1 ± 0.0 | 0.1 ± 0.0 | 0.1 ± 0.0 |
| **Phenylpropionic acids** |  |  |  |  |  |  |  |
| phenylpropionic acid | 0.4 ± 0.1 | 0.0 ± 0.0 | 0.0 ± 0.0 | 0.0 ± 0.0 | 0.0 ± 0.0 | 0.0 ± 0.0 | 0.0 ± 0.0 |
| 2-hydroxyphenylpropionic acid | 0.0 ± 0.0 | 0.0 ± 0.0 | 0.0 ± 0.0 | 0.0 ± 0.0 | 0.0 ± 0.0 | 0.0 ± 0.0 | 0.0 ± 0.0 |
| 3-hydroxyphenylpropionic acid | 0.3 ± 0.0 | 0.0 ± 0.0 | 0.0 ± 0.0 | 0.0 ± 0.0 | 0.0 ± 0.0 | 0.0 ± 0.0 | 0.0 ± 0.0 |
| 4-hydroxyphenylpropionic acid | 0.6 ± 0.0 | 0.0 ± 0.0 | 0.0 ± 0.0 | 0.0 ± 0.0 | 0.0 ± 0.0 | 0.0 ± 0.0 | 0.0 ± 0.0 |
| 4-hydroxy-3-methoxyphenylpropionic acid | 1.7 ± 0.1 | 0.1 ± 0.0 | 0.1 ± 0.1 | 0.3 ± 0.1 | 0.1 ± 0.0 | 0.1 ± 0.0 | 0.2 ± 0.0 |
| **Benzenes** |  |  |  |  |  |  |  |
| 1,2-hydroxybenzene | 4.4 ± 0.4 | 1.0 ± 0.0 | 1.0 ± 0.0 | 2.0 ± 0.0 | 0.9 ± 0.0 | 1.0 ± 0.0 | 1.9 ± 0.0 |
| 1,2,3-trihydroxybenzene | 0.0 ± 0.0 | 0.4 ± 0.0 | 0.4 ± 0.0 | 0.8 ± 0.0 | 0.3 ± 0.0 | 0.3 ± 0.0 | 0.7 ± 0.0 |
| **Acetophenones** |  |  |  |  |  |  |  |
| 4-hydroxyacetophenone | 0.1 ± 0.0 | 0.0 ± 0.0 | 0.0 ± 0.0 | 0.0 ± 0.0 | 0.0 ± 0.0 | 0.0 ± 0.0 | 0.1 ± 0.0 |
| 4-hydroxy-3-methoxyacetophenone | 0.4 ± 0.0 | 0.0 ± 0.0 | 0.0 ± 0.0 | 0.1 ± 0.0 | 0.0 ± 0.0 | 0.0 ± 0.0 | 0.0 ± 0.0 |
| 4-hydroxy-3,5-dimethoxyacetophenone | 0.0 ± 0.0 | 0.0 ± 0.0 | 0.0 ± 0.0 | 0.1 ± 0.0 | 0.0 ± 0.0 | 0.0 ± 0.0 | 0.1 ± 0.0 |
| **Phenylacetic acids** |  |  |  |  |  |  |  |
| phenylacetic acid | 0.4 ± 0.1 | 0.6 ± 0.0 | 0.6 ± 0.0 | 1.2 ± 0.0 | 0.6 ± 0.0 | 0.6 ± 0.0 | 1.2 ± 0.0 |
| 4-hydroxyphenylacetic acid | 4.7 ± 0.4 | 0.3 ± 0.0 | 0.3 ± 0.0 | 0.6 ± 0.0 | 0.3 ± 0.0 | 0.4 ± 0.0 | 0.8 ± 0.0 |
| 3,4-dihydroxyphenylacetic acid | 8.3 ± 4.0. | 0.4 ± 0.1 | 0.5 ± 0.5 | 0.9 ± 0.5 | 2.2 ± 0.0 | 2.3 ± 1.5 | 4.5 ± 1.5 |
| **Mandelic acid** |  |  |  |  |  |  |  |
| 3-hydroxymandelic acid | 0.0 ± 0.0 | 2.8 ± 0.1 | 3.0 ± 0.3 | 5.8 ± 0.3 | 2.8 ± 0.1 | 2.9 ± 0.1 | 5.7 ± 0.2 |
| 4-hydroxymandelic acid | 0.0 ± 0.0 | 25.1 ± 1.8 | 26.0 ± 1.6 | 51.1 ± 2.4 | 22.7 ± 1.2 | 23.1 ± 0.3 | 45.8 ± 1.3 |
| 3,4-dihydroxymandelic acid | 0.0 ± 0.0 | 0.0 ± 0.0 | 0.0 ± 0.0 | 0.0 ± 0.0 | 0.0 ± 0.0 | 0.0 ± 0.0 | 0.0 ± 0.0 |
| **Phenyllactic Acids** |  |  |  |  |  |  |  |
| 4-hydroxyphenyllactic acid | 4.3 ± 0.3 | 2.2 ± 0.0 | 2.6 ± 0.2 | 4.8 ± 0.2 | 2.1 ± 0.0 | 2.4 ± 0.0 | 4.5 ± 0.1 |
| **Phenolics Others** |  |  |  |  |  |  |  |
| chlorogenic acid | 224.3 ± 36.7 | 85.0 ± 2.3 | 85.0 ± 3.7 | 170.0 ± 4.4 | 71.8 ± 2.2 | 71.8 ± 1.5 | 143.7 ± 2.6 |
| ethylferulate | 0.4 ± 0.1 | 0.1 ± 0.0 | 0.1 ± 0.0 | 0.2 ± 0.0 | 0.1 ± 0.0 | 0.1 ± 0.0 | 0.2 ± 0.0 |
| tyrosol | 2.6 ± 0.2 | 1.5 ± 0.0 | 1.7 ± 0.0 | 3.3 ± 0.0 | 1.5 ± 0.1 | 1.7 ± 0.1 | 3.2 ± 0.1 |
| Hydroxytyrosol | 8.3 ± 0.6 | 1.0 ± 0.2 | 1.2 ± 0.0 | 2.2 ± 0.2 | 0.8 ± 0.1 | 0.9 ± 0.0 | 1.7 ± 0.1 |
| **Phenolic dimers** |  |  |  |  |  |  |  |
| Resveratrol | 2.3 ± 0.4 | 0.0 ± 0.0 | 0.0 ± 0.1 | 0.0 ± 0.1 | 0.0 ± 0.1 | 0 ± 0.0 | 0.0 ± 0.1 |
| **Indoles** |  |  |  |  |  |  |  |
| indole-3-acetic acid | 0.2 ± 0.0 | 0.1 ± 0.0 | 0.1 ± 0.0 | 0.2 ± 0.0 | 0.1 ± 0.0 | 0.1 ± 0.0 | 0.2 ± 0.0 |
| indole-3-carboxylic acid | 0.0 ± 0.0 | 0.1 ± 0.0 | 0.1 ± 0.0 | 0.2 ± 0.0 | 0.1 ± 0.0 | 0.1 ± 0.0 | 0.2 ± 0.0 |

Data are presented as mean ± standard deviation or sum ± root mean square, and rounded up.

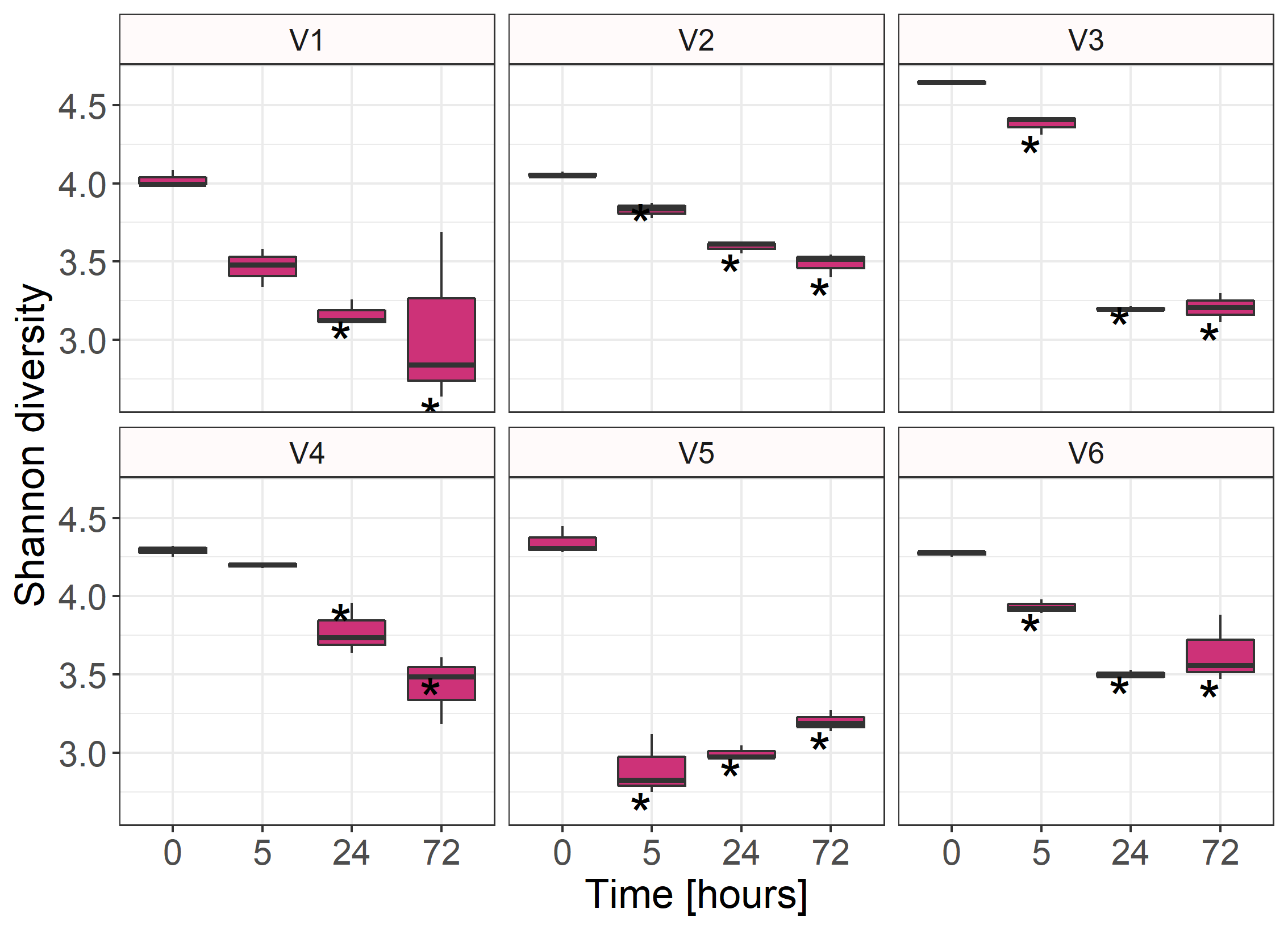

**A**

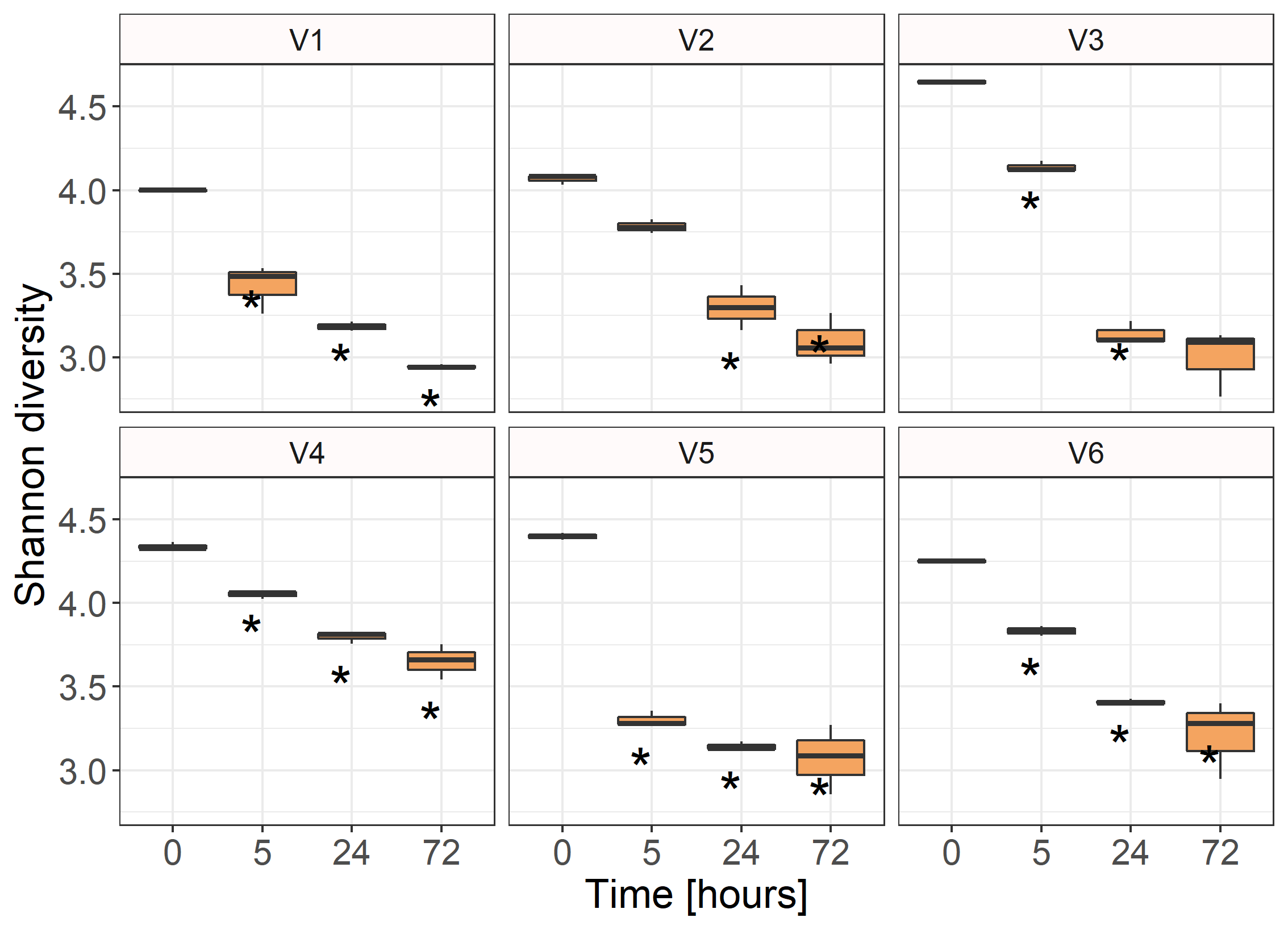

**B**

Supplementary Figure 1: The Shannon diversity index scores for the bacterial communities present in the faecal samples of six healthy donors (V1-V6) over the 72 hour period of the incubation experiments in the presence (purple, **A**) and absence (orange, **B**) of the formulation extract, at baseline (0-hours) and after 5, 24 and 72 hours of incubation. Data are represented as mean ± standard deviation of biological triplicates. Asterisks indicate significantly different donor/sample specific p-values (<0.05), compared to baseline time point.

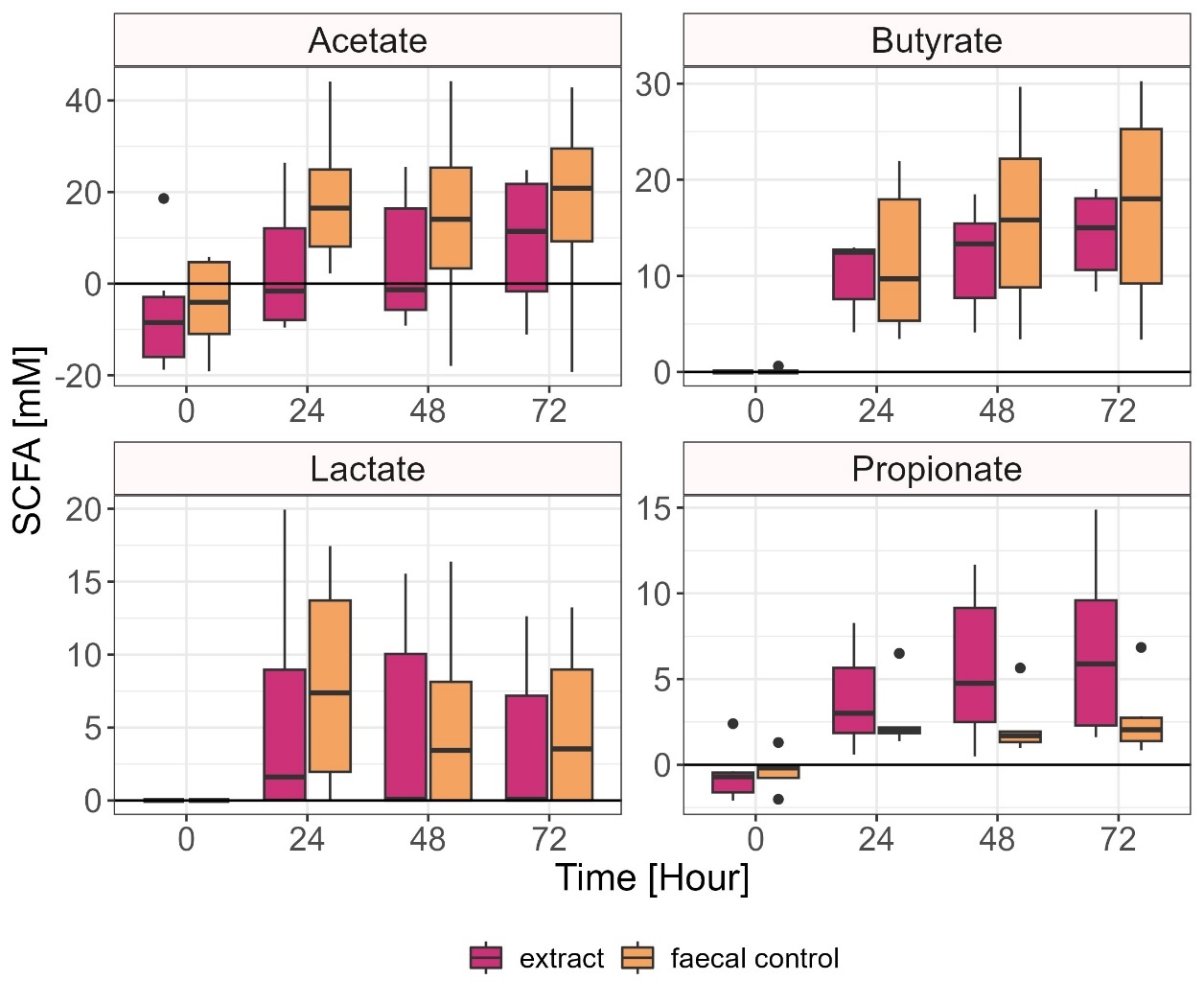

Supplementary Figure 2: Levels of SCFA in samples inoculated with extract (purple) or no extract (orange) at baseline (0 hrs), and after 24, 48 and 72 hrs of inoculation. Levels of SCFA at 0 hours were corrected for levels present in the culturing media. Bars represent mean ± standard deviation from triplicate samples and averaged across six donors. Levels of Acetate were significantly increased at 72hrs compared to baseline in the presence of the extract (p=0.025), and at 24hrs and 72 hrs in the faecal control samples (p=0.019, p=0.032 respectively), when analysing extract and faecal control groups separately. Levels of Butyrate were significantly increased at 24hrs, 48hrs and 72hrs compared to baseline in the presence of the extract (p<0.0001, p<0.0001, p<0.0001 respectively) and in the faecal control samples (p=0.003, p<0.0001, p<0.0001 respectively), when analysing extract and faecal control groups separately. Levels of Propionate were significantly increased at 24hrs, 48hrs and 72hrs compared to baseline in the presence of the extract (p=0.014, p=0.0002, p<0.0001 respectively) and in the faecal control samples (p=0.003, 0.017, 0.003 respectively), when analysing extract and faecal control groups separately. Levels of Lactate were significantly increased at 24hrs in the faecal control samples (p=0.003 respectively), when analysing extract and faecal control groups separately.

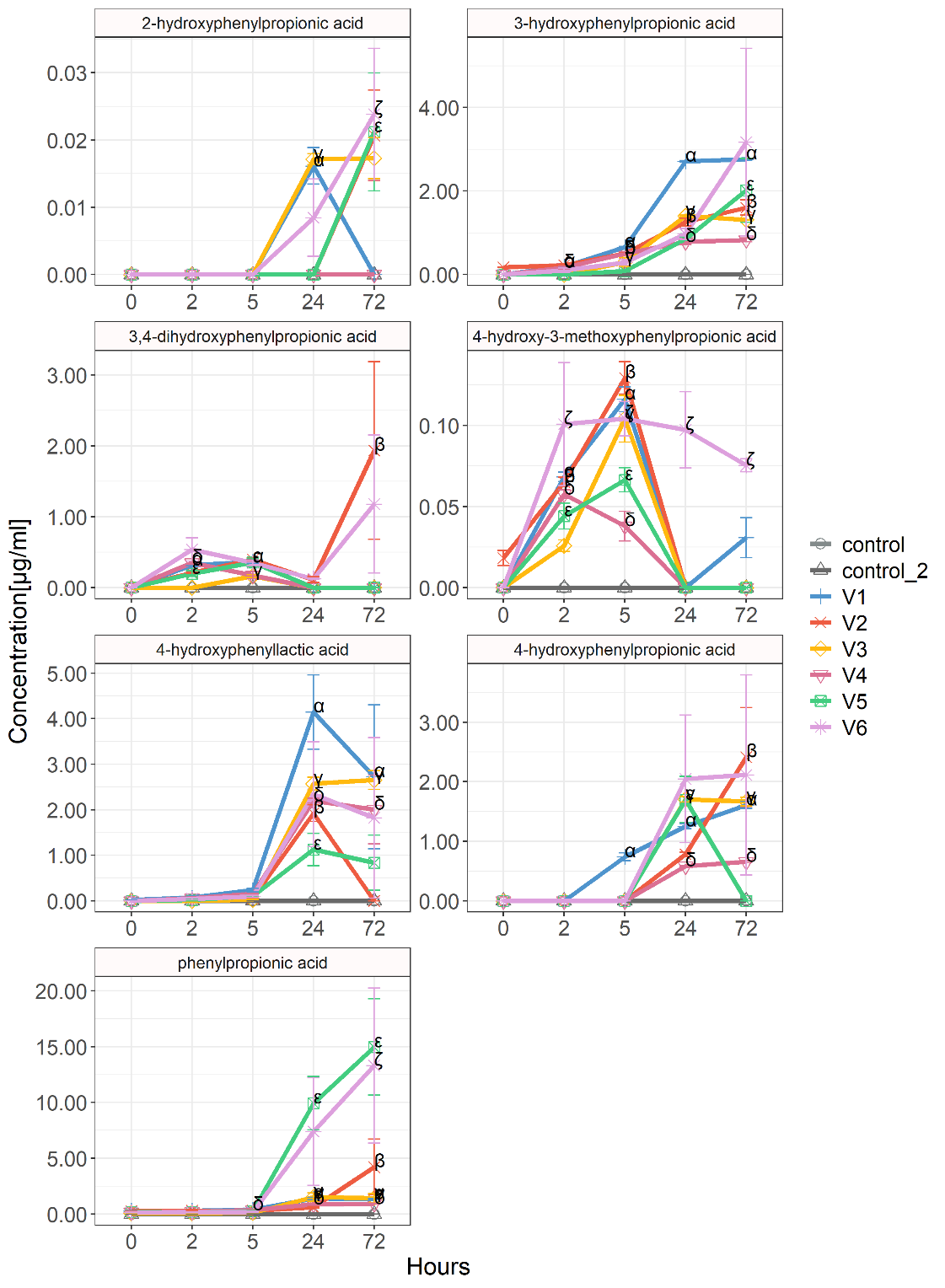

Supplementary Figure 3: Phenylpropionic acid and phenyllactic acid formation by faecal microbiota present in the faecal samples of six healthy donors (V1-V6), and two extract controls (containing only formulation extract and growth media but no faecal bacteria) at baseline (0 hrs), and after 2, 5, 24 to 72 hours of inoculation. donor/sample specific p-values (<0.05): V1-α, V2-β, V3-γ, V4-δ, V5-ε, V6-ζ, control – η, control 2 – θ, compared to the baseline time point.

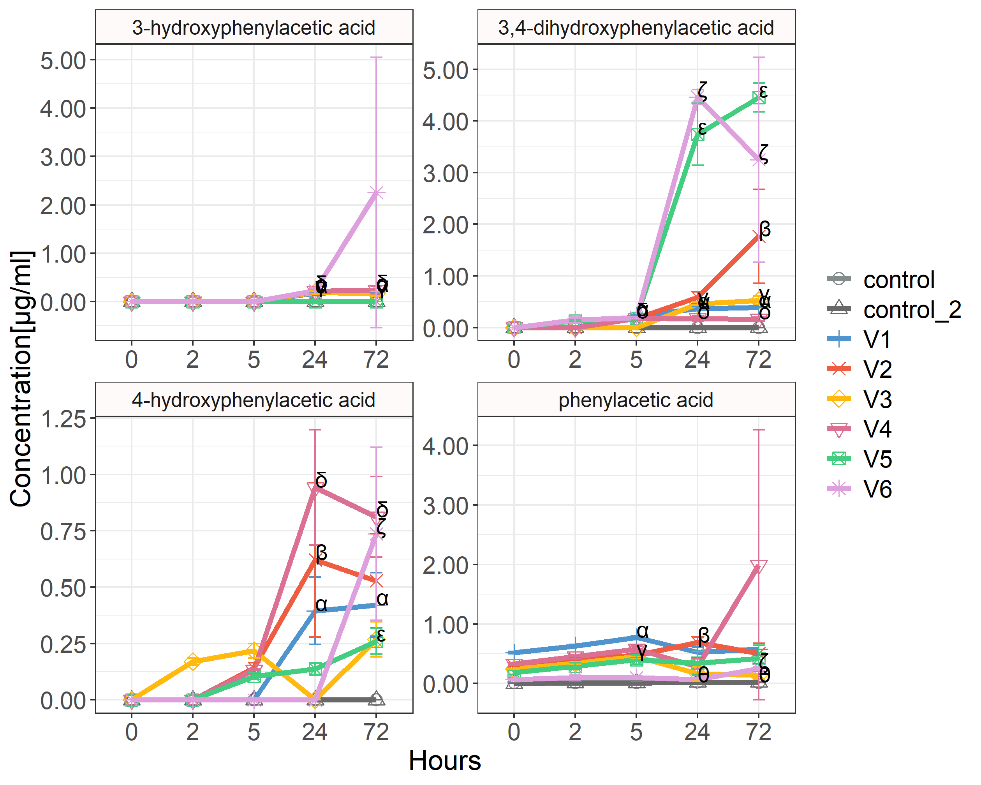

Supplementary Figure 4: Phenylacetic acid formation by faecal microbiota present in the faecal samples of six healthy donors (V1-V6), and two extract controls (containing only formulation extract and growth media but no faecal bacteria) at baseline (0 hrs), and after 2, 5, 24 to 72 hours of inoculation. donor/sample specific p-values (<0.05): V1-α, V2-β, V3-γ, V4-δ, V5-ε, V6-ζ, control – η, control 2 – θ, compared to the baseline time point.

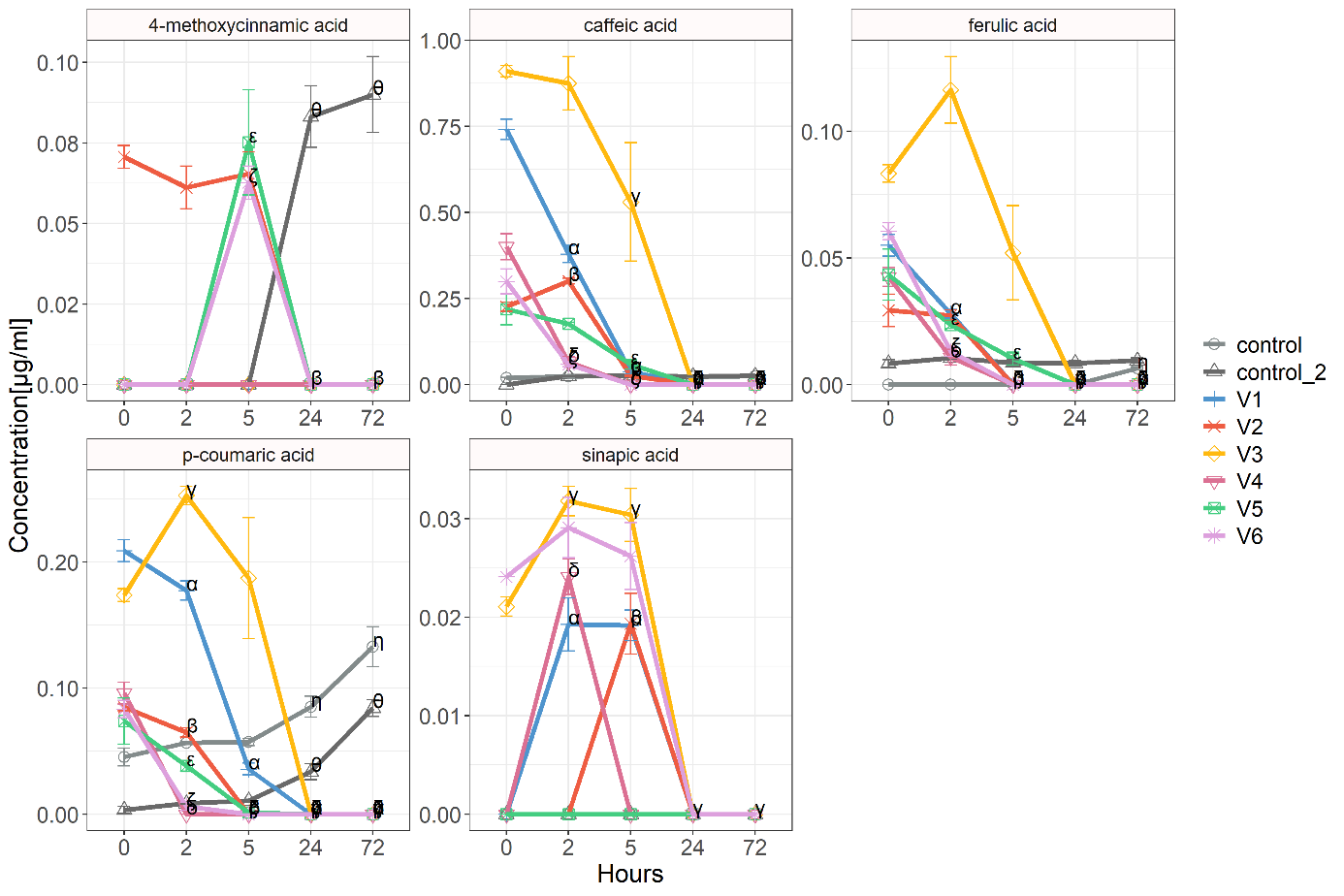

Supplementary Figure 5: Cinnamic acid formation by faecal microbiota present in the faecal samples of six healthy donors (V1-V6), and two extract controls (containing only formulation extract and growth media but no faecal bacteria) at baseline (0 hrs), and after 2, 5, 24 to 72 hours of inoculation. donor/sample specific p-values (<0.05): V1-α, V2-β, V3-γ, V4-δ, V5-ε, V6-ζ, control – η, control 2 – θ, compared to the baseline time point.

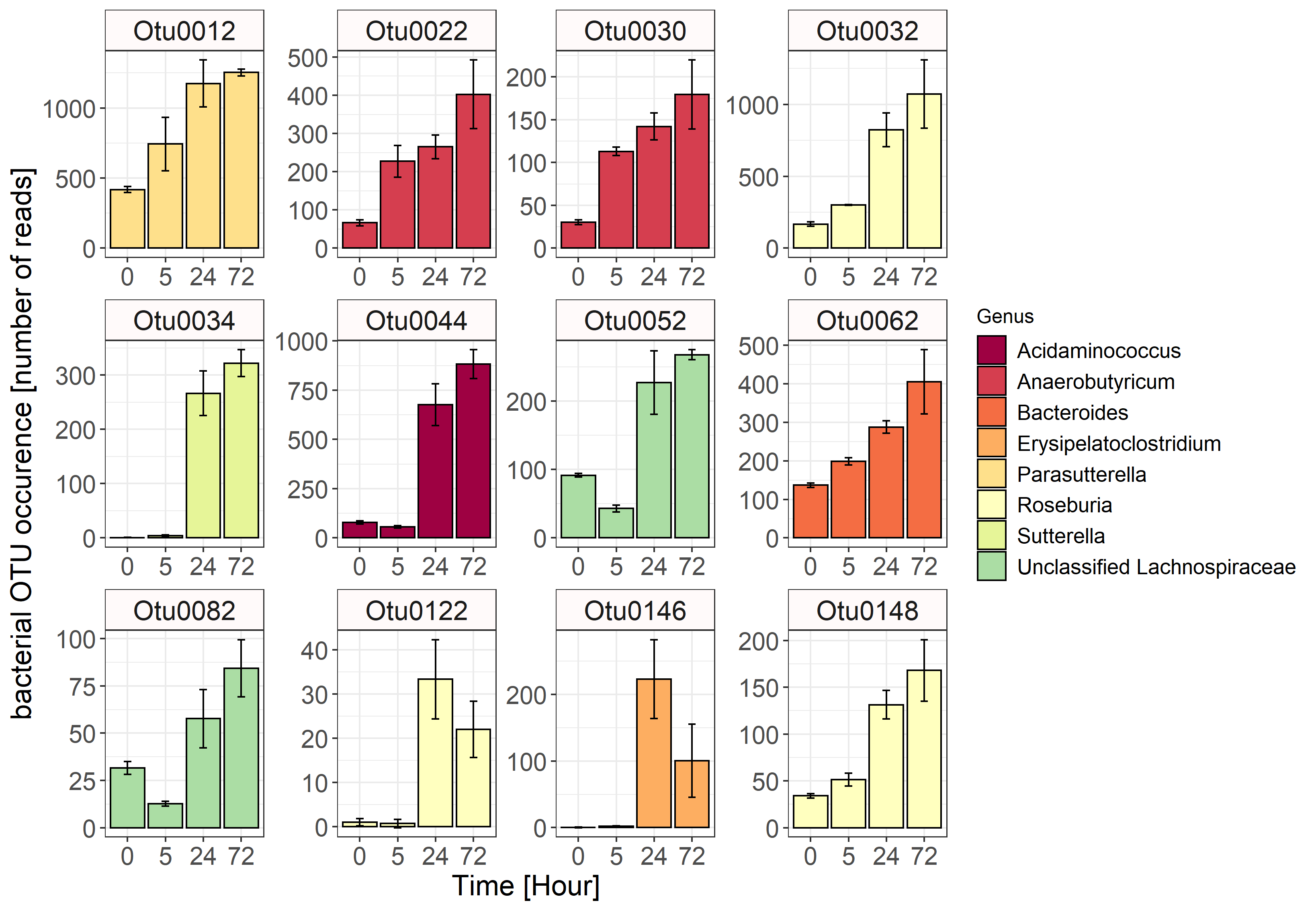

Supplementary Figure 6: Bacterial OTUs from the faecal donor V2, which were negatively correlated with the presence of catechin and epicatechin from the formulation extract. Data are represented as mean ± standard deviation of biological triplicates.

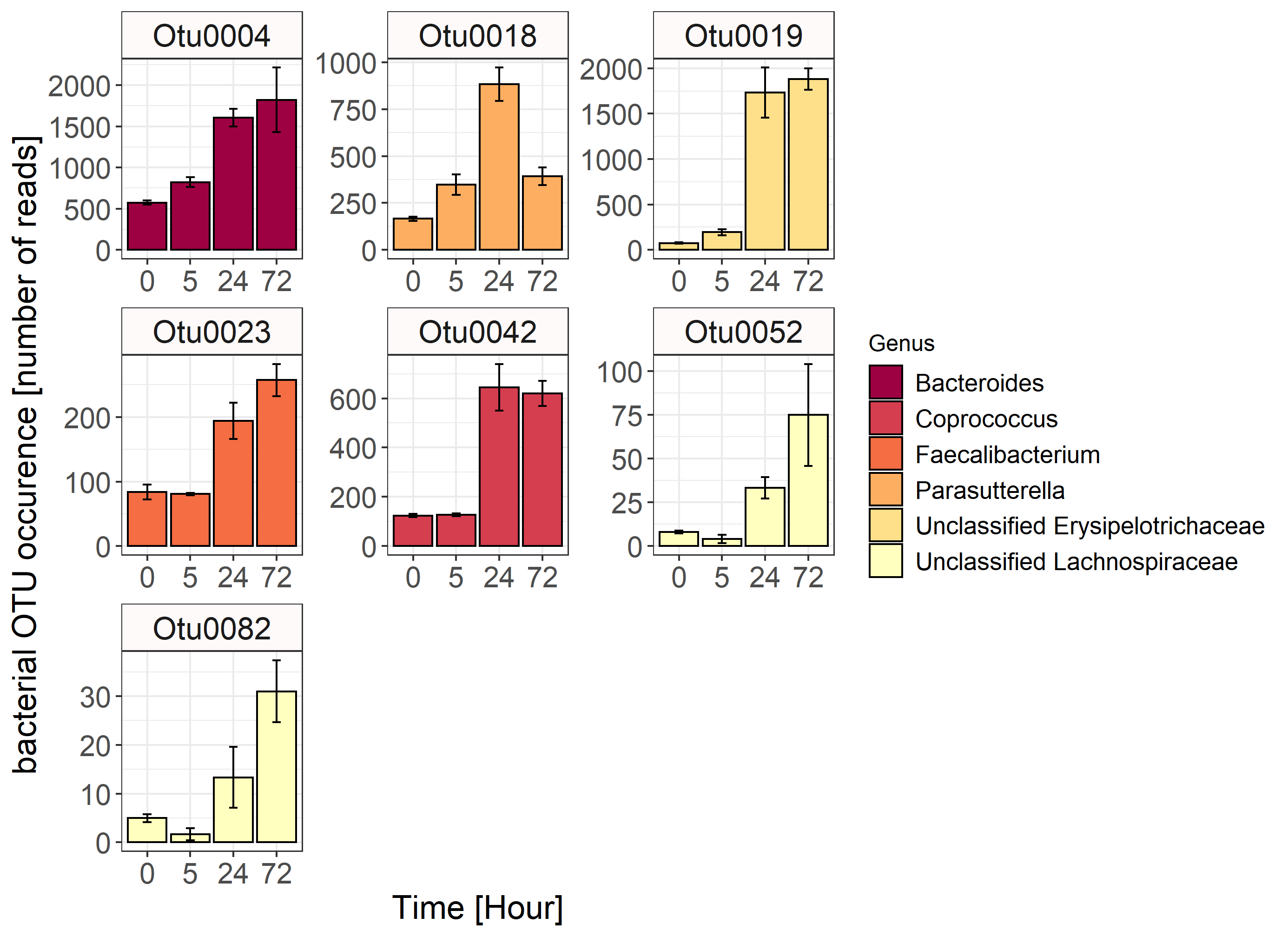

Supplementary Figure 7: Bacterial OTUs from the faecal donor V3, which were negatively correlated with the presence of catechin and epicatechin from the formulation extract. Data are represented as mean ± standard deviation of biological triplicates.

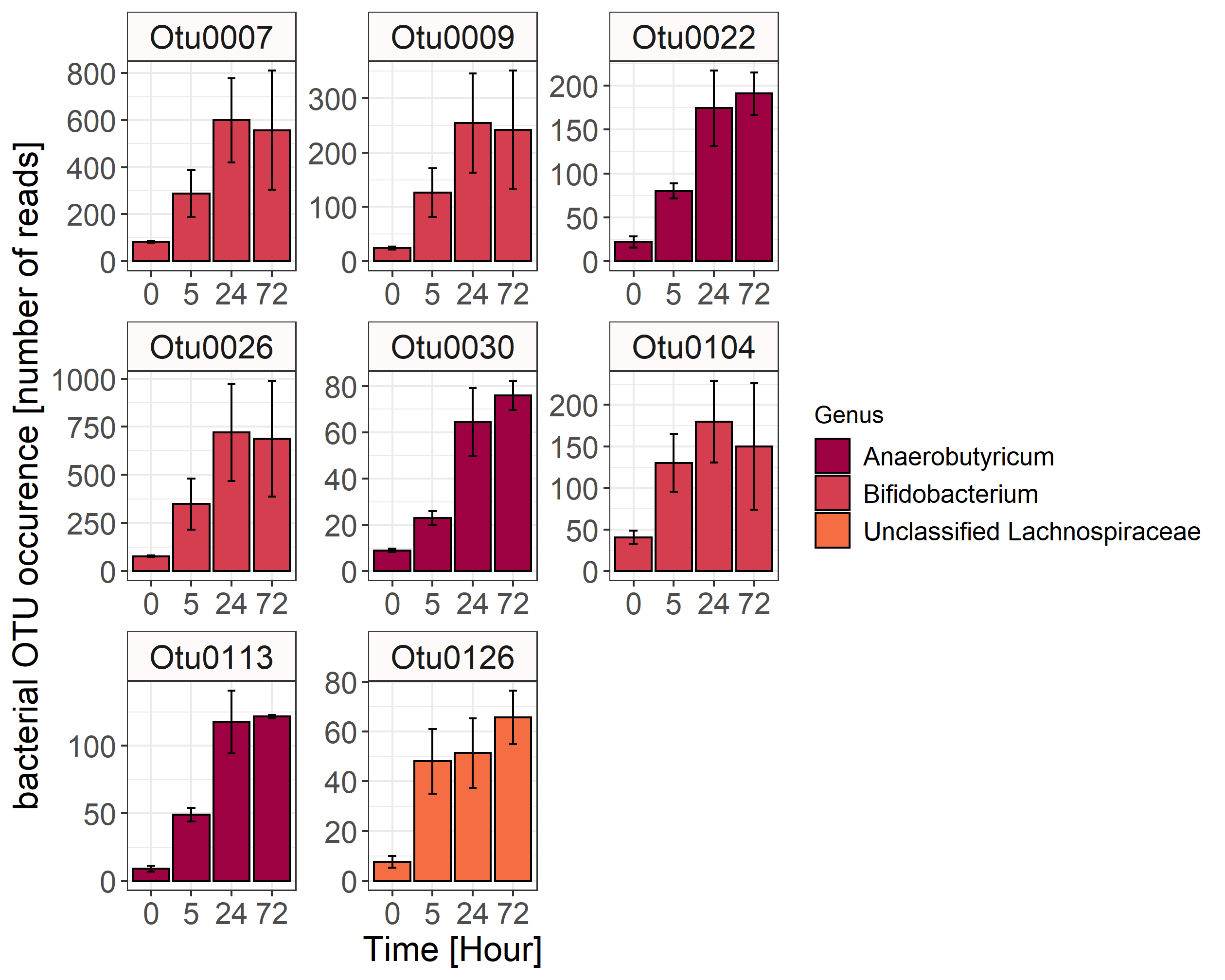

Supplementary Figure 8: Bacterial OTUs from the faecal donor V4, which were negatively correlated with the presence of catechin and epicatechin from the formulation extract. Data are represented as mean ± standard deviation of biological triplicates.

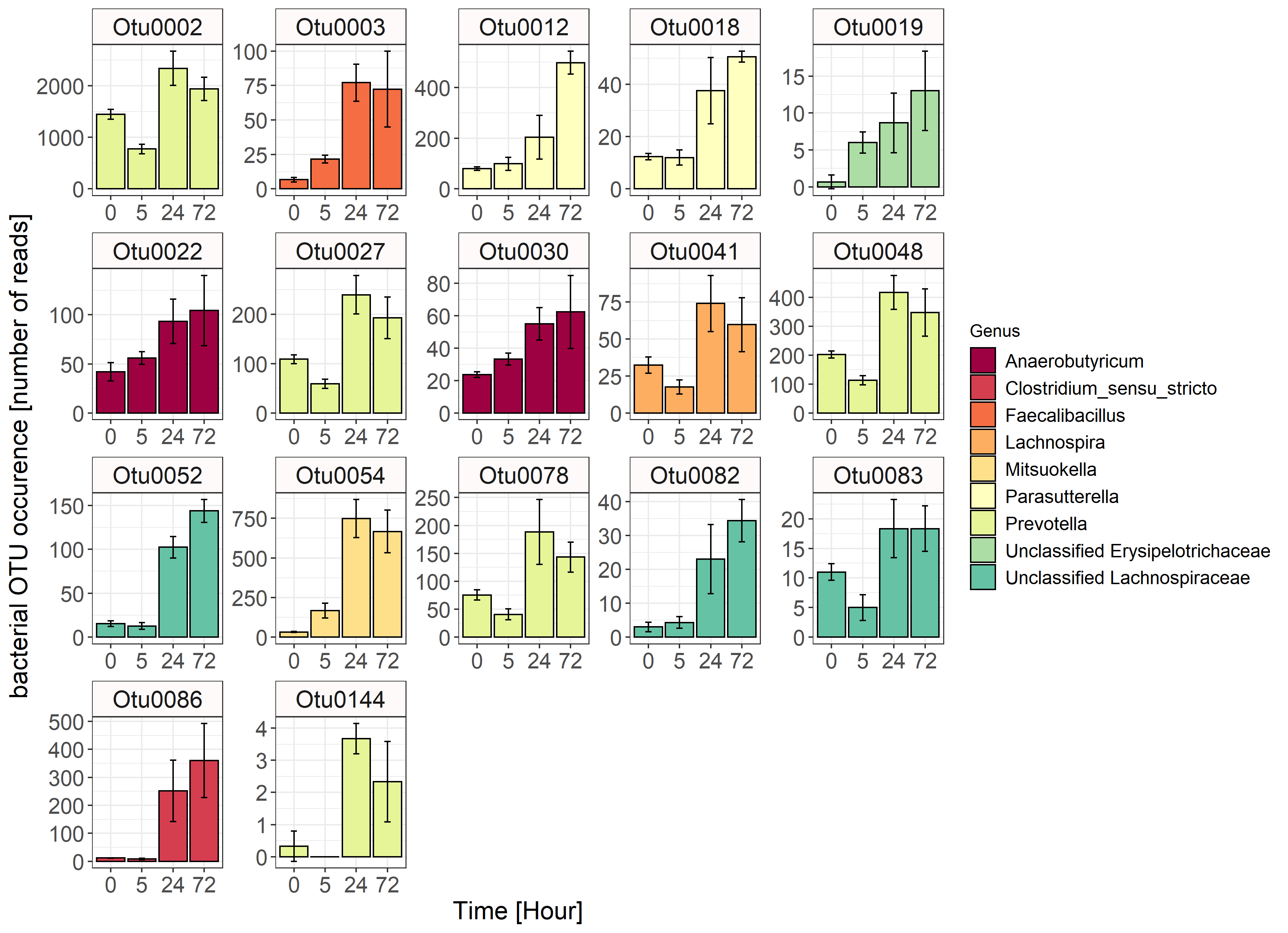

Supplementary Figure 9: Bacterial OTUs from the faecal donor V5, which were negatively correlated with the presence of catechin and epicatechin from the formulation extract. Data are represented as mean ± standard deviation of biological triplicates.

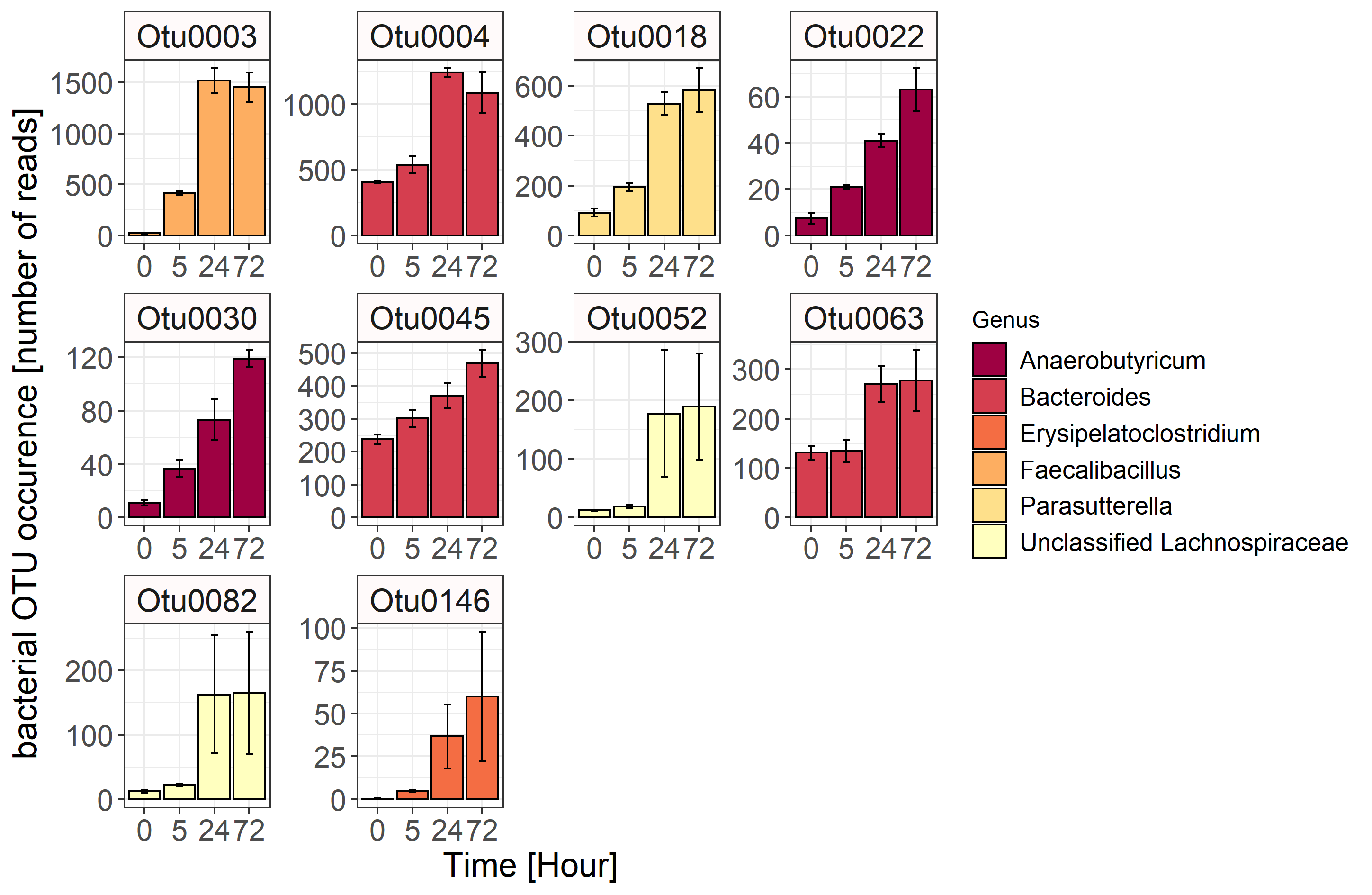

Supplementary Figure 10: Bacterial OTUs from the faecal donor V6, which were negatively correlated with the presence of catechin and epicatechin from the formulation extract. Data are represented as mean ± standard deviation of biological triplicates.

Supplementary Table 6: Identified faecal bacteria that were negatively associated with catechin levels in different Volunteer samples. Numbers in brackets in the Closest BLAST ID column indicate the percentage similarity of hit to the closest cultured species in the NCBI nucleotide reference database. Numbers in brackets after the names in the Genus, Family, Order and Class columns indicate the consistency (out of 100%) of the classifications at each of these taxonomic levels amongst all sequences within a given OTU.

| **V1** | **V2** | **V3** | **V4** | **V5** | **V6** | **Closest BLAST ID** | **Genus (Rep. Seq)** | **Genus (OTU consensus)** | **Family** | **Order** | **Class** |
| --- | --- | --- | --- | --- | --- | --- | --- | --- | --- | --- | --- |
| *Otu0001* | x | x | x | x | x | Catenibacterium mitsuokai (97% similarity) | Catenibacterium(100) | Catenibacterium(100) | Erysipelotrichaceae(100) | Erysipelotrichales(100) | Erysipelotrichia(100) |
| x | x | x | x | *Otu0002* | x | Prevotella copri (99% similarity) | Prevotella(100) | Prevotella(100) | Prevotellaceae(100) | Bacteroidales(100) | Bacteroidia(100) |
| x | x | x | *x* | *Otu0003* | *Otu0003* | Faecalibacillus intestinalis (99% similarity) | Faecalibacillus(100) | Faecalibacillus(100) | Erysipelotrichaceae(100) | Erysipelotrichales(100) | Erysipelotrichia(100) |
| *Otu0004* | x | *Otu0004* | x | x | *Otu0004* | Bacteroides uniformis (98.7% similarity) | Bacteroides(100) | Bacteroides(100) | Bacteroidaceae(100) | Bacteroidales(100) | Bacteroidia(100) |
| x | x | x | *Otu0007* | x | x | Bifidobacterium adolescentis  (96.7% similarity) | Bifidobacterium(100) | Bifidobacterium(100) | Bifidobacteriaceae(100) | Bifidobacteriales(100) | Actinobacteria(100) |
| x | x | x | *Otu0009* | x | x | Bifidobacterium faecale (98.7% similarity) | Bifidobacterium(100) | Bifidobacterium(100) | Bifidobacteriaceae(100) | Bifidobacteriales(100) | Actinobacteria(100) |
| x | *Otu0012* | x | x | *Otu0012* | x | Parasutterella excrementihominis (96.5% similarity) | Parasutterella(100) | Parasutterella(100) | Sutterellaceae(100) | Burkholderiales(100) | Betaproteobacteria(100) |
| *Otu0015* | x | x | x | x | x | Bacteroides dorei (100%) | Bacteroides(100) | Bacteroides(100) | Bacteroidaceae(100) | Bacteroidales(100) | Bacteroidia(100) |
| x | x | *Otu0018* | x | *Otu0018* | *Otu0018* | Parasutterella excrementihominis (99.7% similarity) | Parasutterella(100) | Parasutterella(100) | Sutterellaceae(100) | Burkholderiales(100) | Betaproteobacteria(100) |
| x | x | *Otu0019* | x | *Otu0019* | x | uncultured, Longibaculum sp. strain 2P-4 (90.1% similarity) | Unclassified Erysipelotrichaceae(100) | Unclassified Erysipelotrichaceae(100) | Erysipelotrichaceae(100) | Erysipelotrichales(100) | Erysipelotrichia(100) |
| *Otu0022* | *Otu0022* | x | *Otu0022* | *Otu0022* | *Otu0022* | Anaerobutyricum hallii (98.4% similarity) | Anaerobutyricum(100) | Anaerobutyricum(100) | Lachnospiraceae(100) | Clostridiales(100) | Clostridia(100) |
| *Otu0023* | x | *Otu0023* | x | x | x | Faecalibacterium prausnitzii (96.1% similarity) | Faecalibacterium(100) | Faecalibacterium(100) | Ruminococcaceae(100) | Clostridiales(100) | Clostridia(100) |
| *Otu0024* | x | x | x | x | x | Bacteroides ovatus (99.7% similarity) | Bacteroides(100) | Bacteroides(100) | Bacteroidaceae(100) | Bacteroidales(100) | Bacteroidia(100) |
| x | x | x | *Otu0026* | x | x | Bifidobacterium faecale (98% similarity) | Bifidobacterium(100) | Bifidobacterium(100) | Bifidobacteriaceae(100) | Bifidobacteriales(100) | Actinobacteria(100) |
| x | x | x | x | *Otu0027* | x | Prevotella copri (97.2% similarity) | Prevotella(96) | Prevotella(100) | Prevotellaceae(100) | Bacteroidales(100) | Bacteroidia(100) |
| *Otu0030* | *Otu0030* | x | *Otu0030* | *Otu0030* | *Otu0030* | Anaerobutyricum hallii (96.9% similarity) | Anaerobutyricum(100) | Anaerobutyricum(100) | Lachnospiraceae(100) | Clostridiales(100) | Clostridia(100) |
| *Otu0031* | x | x | x | x | x | Bacteroides fragilis (100%) | Bacteroides(100) | Bacteroides(100) | Bacteroidaceae(100) | Bacteroidales(100) | Bacteroidia(100) |
| *Otu0032* | *Otu0032* | x | x | *x* | x | Roseburia intestinalis (100%) | Roseburia(100) | Roseburia(100) | Lachnospiraceae(100) | Clostridiales(100) | Clostridia(100) |
| x | *Otu0034* | x | x | x | x | Sutterella wadsworthensis (100%) | Sutterella(100) | Sutterella(100) | Sutterellaceae(100) | Burkholderiales(100) | Betaproteobacteria(100) |
| x | x | x | x | *Otu0041* | x | Lachnospira eligens (100%) | Lachnospira(100) | Lachnospira(100) | Lachnospiraceae(100) | Clostridiales(100) | Clostridia(100) |
| x | x | *Otu0042* | x | x | x | Coprococcus eutactus (99% similarity) | Coprococcus(99) | Coprococcus(100) | Lachnospiraceae(100) | Clostridiales(100) | Clostridia(100) |
| x | *Otu0044* | x | x | x | x | Acidaminococcus intestini (99.7% similarity) | Acidaminococcus(100) | Acidaminococcus(100) | Acidaminococcaceae(100) | Selenomonadales(100) | Negativicutes(100) |
| x | x | x | x | x | *Otu0045* | Bacteroides thetaiotaomicron (100%) | Bacteroides(100) | Bacteroides(100) | Bacteroidaceae(100) | Bacteroidales(100) | Bacteroidia(100) |
| x | x | x | x | *Otu0048* | x | Prevotella copri (97.2% similarity) | Prevotella(94) | Prevotella(100) | Prevotellaceae(100) | Bacteroidales(100) | Bacteroidia(100) |
| *Otu0050* | x | x | x | x | x | Lactobacillus mucosae (100%) | Lactobacillus(100) | Lactobacillus(100) | Lactobacillaceae(100) | Lactobacillales(100) | Bacilli(100) |
| x | *Otu0052* | *Otu0052* | x | *Otu0052* | *Otu0052* | Enterocloster bolteae (96% similarity) | Unclassified Lachnospiraceae(81) | Unclassified Lachnospiraceae(100) | Lachnospiraceae(100) | Clostridiales(100) | Clostridia(100) |
| x | x | x | x | *Otu0054* | x | Mitsuokella jalaludinii (99% similarity) | Mitsuokella(100) | Mitsuokella(100) | Veillonellaceae(100) | Selenomonadales(100) | Negativicutes(100) |
| *Otu0061* | x | x | x | x | x | Bacteroides xylanisolvens (100%) | Bacteroides(100) | Bacteroides(100) | Bacteroidaceae(100) | Bacteroidales(100) | Bacteroidia(100) |
| x | *Otu0062* | x | x | x | x | Bacteroides faecis (100%) | Bacteroides(100) | Bacteroides(100) | Bacteroidaceae(100) | Bacteroidales(100) | Bacteroidia(100) |
| x | x | x | x | x | *Otu0063* | Bacteroides oleiciplenus (98.4% similarity) | Bacteroides(100) | Bacteroides(100) | Bacteroidaceae(100) | Bacteroidales(100) | Bacteroidia(100) |
| *Otu0067* | x | x | x | x | x | Bacteroides caccae (99.7% similarity) | Bacteroides(100) | Bacteroides(100) | Bacteroidaceae(100) | Bacteroidales(100) | Bacteroidia(100) |
| x | x | x | x | *Otu0078* | x | Prevotellaceae copri (95.6% similarity) | Prevotella(98) | Prevotella(100) | Prevotellaceae(100) | Bacteroidales(100) | Bacteroidia(100) |
| *Otu0081* | x | x | x | x | x | Duodenibacillus massiliensis (100%) | Parasutterella(65) | Unclassified Burkholderiales(99) | Unclassified Burkholderiales(99) | Burkholderiales(100) | Betaproteobacteria(100) |
| x | *Otu0082* | *Otu0082* | x | *Otu0082* | *Otu0082* | Enterocloster bolteae (96% similarity) | Unclassified Lachnospiraceae(93) | Unclassified Lachnospiraceae(100) | Lachnospiraceae(100) | Clostridiales(100) | Clostridia(100) |
| *Otu0083* | x | x | x | *Otu0083* | x | [Lactobacillus] rogosae (99.7% similarity) | Unclassified Lachnospiraceae(100) | Unclassified Lachnospiraceae(100) | Lachnospiraceae(100) | Clostridiales(100) | Clostridia(100) |
| x | x | *x* | x | *Otu0086* | x | Clostridium vincentii (96.7% similarity) | Clostridium_sensu_stricto(100) | Clostridium_sensu_stricto(100) | Clostridiaceae_1(100) | Clostridiales(100) | Clostridia(100) |
| x | x | x | *Otu0104* | x | x | Bifidobacterium pseudocatenulatum (97%) | Bifidobacterium(100) | Bifidobacterium(100) | Bifidobacteriaceae(100) | Bifidobacteriales(100) | Actinobacteria(100) |
| x | x | x | *Otu0113* | x | x | Anaerobutyricum hallii (96.9% similarity) | Anaerobutyricum(100) | Anaerobutyricum(100) | Lachnospiraceae(100) | Clostridiales(100) | Clostridia(100) |
| *Otu0122* | *Otu0122* | x | x | x | x | Roseburia hominis (100%) | Roseburia(100) | Roseburia(100) | Lachnospiraceae(100) | Clostridiales(100) | Clostridia(100) |
| x | x | x | *Otu0126* | x | x | Coprococcus comes (97.2% similarity) | Unclassified Lachnospiraceae(100) | Unclassified Lachnospiraceae(100) | Lachnospiraceae(100) | Clostridiales(100) | Clostridia(100) |
| *Otu0140* | x | x | x | *x* | x | [Lactobacillus] rogosae (97.9% similarity) | Unclassified Lachnospiraceae(100) | Unclassified Lachnospiraceae(100) | Lachnospiraceae(100) | Clostridiales(100) | Clostridia(100) |
| x | x | x | x | *Otu0144* | x | Prevotella copri (95.3% similarity) | Prevotella(96) | Prevotella(100) | Prevotellaceae(100) | Bacteroidales(100) | Bacteroidia(100) |
| x | *Otu0146* | x | x | x | *Otu0146* | Erysipelatoclostridium ramosum (100%) | Erysipelatoclostridium(100) | Erysipelatoclostridium(100) | Erysipelotrichaceae(100) | Erysipelotrichales(100) | Erysipelotrichia(100) |
| *Otu0147* | x | x | x | x | x | Bacteroides stercoris (99.4% similarity) | Bacteroides(100) | Bacteroides(100) | Bacteroidaceae(100) | Bacteroidales(100) | Bacteroidia(100) |
| x | *Otu0148* | x | x | x | x | Roseburia intestinalis (96.7% similarity) | Roseburia(97) | Roseburia(100) | Lachnospiraceae(100) | Clostridiales(100) | Clostridia(100) |

Supplementary Table 7: Levels of Epicatechin and Epigallocatechin at timepoint 0 and 24 hrs in the presence of single bacterial cultures and formulation extract.

|  | **Epicatechin** | | **Epigallocatechin** | |
| --- | --- | --- | --- | --- |
|  | **0 hours** | **24 hours** | **0 hours** | **24 hours** |
| *Anaerobutyricum hallii DSM3353* | 1133.6 ± 124.1 | 1492.4 ± 160.7 | 18.5 ± 1.7 | 27.7 ± 2.0 |
| *Anearobutyricum soehngenii L2-7* | 1533.1 ± 212.6 | 1562.5 ± 219.8 | 68.2 ± 7.1 | 64.5 ± 3.9 |
| *Enterocloster bolteae R7* | 1305.6 ± 135.6 | 1457.2 ± 166.8 | 55.7 ± 1.9 | 73.1 ± 3.0 |
| *Fusicatenibacter saccharivorans D5 BHI MAN 8* | 1324.7 ± 107.2 | 1548.8 ± 117.1 | 53.0 ± 1.6 | 75.0 ± 3.7 |
| *Coprococcus* sp*.* L2-50 | 1394.7 ± 170.9 | 1477.8 ± 194.0 | 26.3 ± 21.7 | 57.6 ± 3.2 |
| *Flavonifractor plautii DSM4000* | 1277.1 ± 100.8 | 1503.5 ± 83.1 | 0±0 | 54.6 ± 0.7 |
| Control | 1573.5 | 1665.1 | 0.0 | 0.0 |

Data are presented as mean ± standard deviation of triplicate cultures. Control samples were tested in singlet. Data were analysed via T-test comparing individual flavonoid concentrations in each single bacterial strain between 0 and 24 hours. * - significant p-value <0.05.
